# Gene expression and chromatin accessibility during progressive EMT and MET linked to dynamic CTCF engagement

**DOI:** 10.1101/2020.05.11.089110

**Authors:** Kelsey S. Johnson, Shaimaa Hussein, Shuxuan Song, Priyanka Chakraborty, Mohit Kumar Jolly, Michael J. Toneff, Yin C. Lin, Joseph H. Taube

## Abstract

**Background:** Epithelial-mesenchymal transition (EMT) facilitates cellular movements critical for proper development; however, in a carcinoma, EMT promotes metastatic dissemination. Stable intermediate states (partial-EMT) are increasingly implicated in metastatic dissemination while reversal of EMT, termed mesenchymal-epithelial transition (MET), is increasingly implicated in metastatic colonization. To understand the partial and reversible nature of EMT, we characterized chromatin accessibility dynamics, transcriptome changes, protein expression patterns, as well as E-cadherin expression, localization, and gene-level dynamics in mammary epithelial cells undergoing stepwise reversible EMT.

**Results:** While shorter EMT induction induced internalization of E-cadherin protein, surface expression was recovered upon MET without loss of transcript or bulk protein. Conversely, a longer EMT induced stable repression of E-cadherin indicated by loss of chromatin accessibility and induced global expansion of accessible sites across the genome, facilitated by increased engagement of multiple transcription factor families, including AP-1 and SMAD. We observe enrichment for binding sites for the insulator proteins CTCF and BORIS was significantly diminished in both stemness-enriched partial-EMT and partial-MET states and determined that CTCF repression imparts alterations in some histone covalent modifications concomitant with those observed during TGFβ-induced EMT.

**Conclusions:** These findings are indicative of a major role for chromatin looping and reorganization in plasticity, stemness, and partial EMT phenotypes.

## Background

Epithelial-mesenchymal transition (EMT) is a conserved cellular process that drives programs such as gastrulation and wound healing. During EMT, epithelial cells alter their gene expression and morphology, lose cell-cell contacts, and adopt a mesenchymal-like state (1). Because this process promotes invasion, intravasation, and resistance to anoikis in tumor cells, the EMT program is also implicated in metastatic dissemination (2-4). Recent work has contributed to a revised model of metastasis in which reversal of EMT, mesenchymal-epithelial transition (MET), is necessary for colonization of cells which arrive at the metastatic site by means of an EMT (5-8). Thus, understanding the factors that impact the reversibility of EMT is critical to developing better anti-metastasis treatments.

EMT propels cells through a progressive adoption of gene expression changes leading to phenotypic alterations. While a hallmark of EMT is the suppression of genes such as E-cadherin (*CDH1*) and epithelial cell adhesion molecule (*EPCAM*), there are also profound changes throughout the epigenome, transcriptome, spliceosome, and protein translation machinery (9). EMT can be initiated by microenvironmental signals such as TGFβ, EGF, hypoxia, and tissue stiffness (10-13), and is effected through networks of EMT-transcription factor proteins (EMT-TFs) such as SNAIL, SLUG, TWIST1, ZEB1, SIX1, SOX10, and FOXC2 (14-18). These transcription regulators act in conjunction with epigenetic regulatory mechanisms such as histone post-translational modifications, re-organized large organized heterochromatin K9 modifications (LOCK) domains (19-22), DNA-methylation (21, 23, 24), and microRNAs including the miR-200 family (25-28) and miR-203 (29, 30) among others.

Conversely MET stimulates the re-emergence of the epithelial phenotype, re-establishment of cell-cell contacts, a decrease in migratory traits, and re-expression of transcription factors such as ELF5, GRHL2, and OVOL1/2 (31-33).

Despite considerable overlap within the EMT-regulatory network, exogenous expression of individual EMT-TFs leads to distinctive EMT states (15, 34). Moreover, diverse partial- or hybrid-EMT, which express epithelial and mesenchymal traits have been recently described (35-40) and are reported to be highly plastic, efficiently initiate tumor growth (37), and indicate to poorer patient outcome (18, 40-44). Nevertheless, a multi-level analysis, including assessment of chromatin accessibility of cells experimentally induced to undergo EMT *and* MET has not been shown.

We endeavored to determine the genome-wide dynamics of chromatin accessibility at multiple timepoints during EMT and MET and characterize the relationship between chromatin accessibility, gene expression, and cell phenotype. Utilizing the <u>a</u>ssay for transposase-<u>a</u>ccessible <u>c</u>hromatin with next-generation <u>seq</u>uencing (ATAC-seq), we report that MCF10A mammary epithelial cells, proceeding through stepwise EMT and MET, undergo progressive and semi-reversible alterations in chromatin accessibility. By staggering the time of TGFβ exposure and removal, we establish a series of partial states exhibiting distinct E-cadherin gene, transcript, and protein dynamics. Our analyses of chromatin accessibility in these states reveal that progression through EMT imparts a global increase in chromatin accessibility while MET is marked by chromatin compaction. We show that the chromatin insulator protein CTCF is a major participant in oscillating chromatin dynamics between EMT-induced and MET-resolved states and is required for preventing EMT-induced accumulation of the LOCK-domain associated mark, H3K9me3. Importantly, we find that initiation of EMT has the capacity to instill durable effects on the chromatin structure--most notably loss of accessibility at the E-cadherin promoter despite MET, potentially allowing the detection of past EMT events. Collectively our findings indicate that activation of EMT and MET dramatically reconfigures chromatin organization with distinct effects dependent on the duration of EMT-inducing signal.

## Results

### TGFβ induced morphological and transcriptomic dynamics

MCF10A human epithelial cells derived from spontaneously immortalized fibrocystic mammary tissue (45) have long been used as a model for epithelial-mesenchymal plasticity (46-49). To characterize progressive EMT/MET, we exposed cells to TGFβ treatment and withdrawal for varying durations (50). The characteristic epithelial was lost at 2 days of TGFβ treatment while typical spindle-like morphology emerged at 4 days of treatment (Fig. 1a). Notably, 4 days of TGFβ withdrawal following a short-term treatment (4 days) elicited a rapid return to epithelial morphology (Fig. 1a) while 10 days of withdrawal was necessary to observe a return to epithelial morphology following long-term (10 days) treatment (Fig. 1a). To gauge the effect of varying durations of exposure to EMT-inducers, we continued to use both short-term treatment (4 days of TGFβ, then withdrawal) and long-term treatment (10 days of TGFβ, then withdrawal) models.

**Fig. 1.**
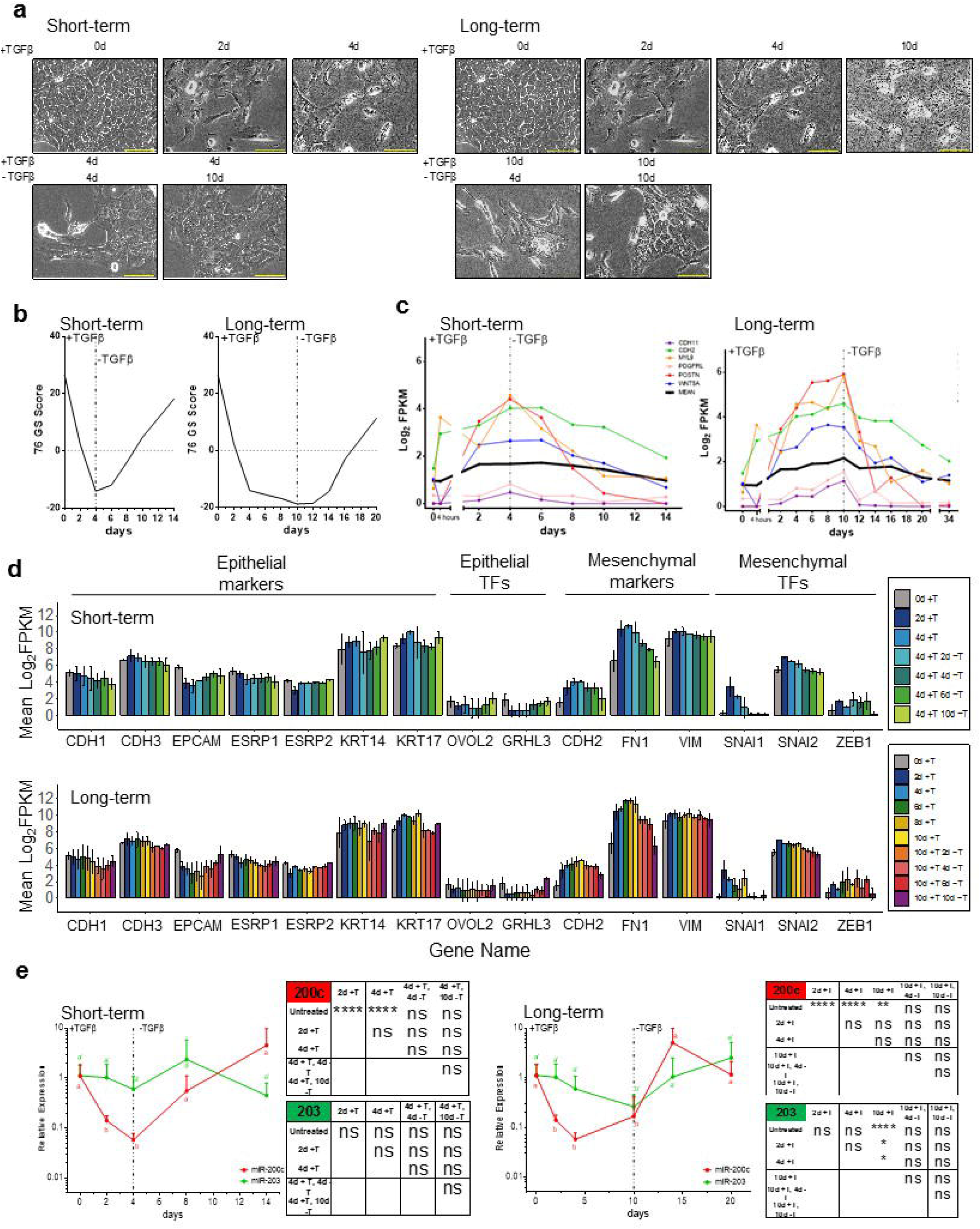
Phased gene expression changes during TGFβ treatment and withdrawal. **a** Brightfield photomicrographs of MCF10A cells at indicated durations of 5 ng/mL TGFβ treatment and withdrawal in short-term (left) and long-term (right) TGFβ-induced EMT models. Scale bars = 80 µm. mRNA expression patterns based on the: **b** 76-Gene Epithelial Metric (51), and **c** log_2_FPKM expression of Core EMT Signature (34) in matched replicate (n = 1). **d** mRNA expression of select epithelial and mesenchymal genes as defined by RNA-seq in short-term (top) and long-term (bottom) models (n = 2, mean ± SEM). **e** Expression of miR-200c (red) and miR-203 (green) during short-term (left) and long-term (right) models. Data were normalized using sno-U6 and shown as mean + SEM from n = 4 with the indicated significance by using a two-tailed Student’s *t* test, statistical comparisons shown in table to the right of graph. ns = not significant, \**p* ≤ 0.05, \*\**p* ≤ 0.01, **** *<u>p</u>* < 0.0001

We next assessed changes in gene expression by RNA-seq. To measure broad changes in epithelial gene expression, we applied an epithelial-specific 76-gene score (76GS), where higher scores correspond to an epithelial gene expression pattern (51). TGFβ treatment suppressed the 76GS score beginning at 2 days of treatment until 2 days following withdrawal whereupon it slowly recovered with additional withdrawal time (Fig. 1b). This analysis was followed by segmentation using a mammary cell-specific EMT gene expression signature (34). Gene expression data was selected for 50 genes universally upregulated by distinct models of EMT in HMLE mammary cells (34). In our model, EMT-Up genes increased expression at different timepoints, indicating a phased implementation of gene activation (Fig. 1c). Excepting *CDH2*, the majority of EMT-Up core genes were rapidly suppressed. Withdrawal in from short-term treatment also facilitated a return to untreated levels of gene expression in most genes (Fig. 1c).

When examining regulation of individual genes, TGFβ expectedly downregulated expression of epithelial-related genes such as *CDH1, EPCAM, OVOL2*, and *GRHL3* while withdrawal increased expression of these genes (Fig. 1d). *CDH3* (P-cadherin) a proposed partial-EMT marker (52) followed a similar pattern as *CDH1* (Fig 1d). We also characterized gene expression patterns of mesenchymal-related genes *CDH2, FN1, SNAI1/2, VIM, and ZEB1*, and observed upregulation following TGFβ treatment and downregulation following withdrawal in both models (Fig. 1d).

Given the modest changes in the steady-state levels of *CDH1* mRNA, we profiled the dynamics of *CDH1* transcription regulation using a promoter reporter. MCF10A cells were transduced with the *CDH1* promoter linked to an RFP-encoding gene, facilitating promoter activity tracking at the single cell level (23). Commensurate with RNA-seq data, the *CDH1* promoter reporter remained active throughout the short-term TGFβ treatment and withdrawal timecourse (Sup. Fig. 1a,b, Tables 1,2 pink panel). However, upon extended treatment, reporter-negative cells outnumbered reporter-positive cells, indicating promoter-induced repression (Sup. Fig. 1a,b, Tables 1,2 pink panel). Notably, the repression of the *CDH1* promoter was durable despite ten days of withdrawal.

We extended our observation of transcriptional dynamics during reversible EMT to microRNAs 200c and 203, which have been reported to target *ZEB1/2* mRNA (25, 53). In both models, miR-200c and miR-203 decrease expression following 4 days of TGFβ treatment and 10 days of TGFβ treatment results in stronger suppression for both miRNAs (Fig 1e).

Our data demonstrate that individual gene expression dynamics follow distinct kinetics during reversible EMT. In particular, our model exhibited diminished total epithelial gene expression early during TGFβ treatment, pointing to a partial-EMT state characterized by a modest decrease in *EPCAM* and *CDH1*, upregulation of *SNAI1/2*, and rapid activation of signaling effectors such as *POSTN* and *WNT5A*. During MET, a long-term TGFβ-driven EMT is distinguished from short-term TGFβ-driven EMT by a failure to re-activate *CDH1* promoter activity. Thus, within tumors, cells that do not express *CDH1* may be nevertheless undergoing MET.

### TGFβ induced protein dynamics

Many EMT-related genes are regulated by translation or protein localization (48, 54). Consequently, we assessed protein levels of EMT markers and EMT-TFs. We observed that TGFβ induced suppression of E-cadherin and increased expression of mesenchymal markers N-cadherin and vimentin (Fig. 2a). In our short-term model, TGFβ withdrawal led to protein expression typical of untreated cells, while long-term TGFβ treatment led to more durable changes following withdrawal (Fig. 2a). For example, E-cadherin expression remained suppressed through 4 days of withdrawal and N-cadherin expression remained elevated through 10 days of withdrawal (Fig. 2a). Expression of the EMT-TF Slug increased within 4 hours of initiating TGFβ treatment while expression of EMT-TFs Twist and ZEB1 did not emerge until later in the ong-term TGFβ treatment model (Fig. 2a), indicative of a progressive activation of EMT-TFs wherein partial-EMT is driven by Slug prior to expression of Twist and ZEB1.

**Fig. 2.**
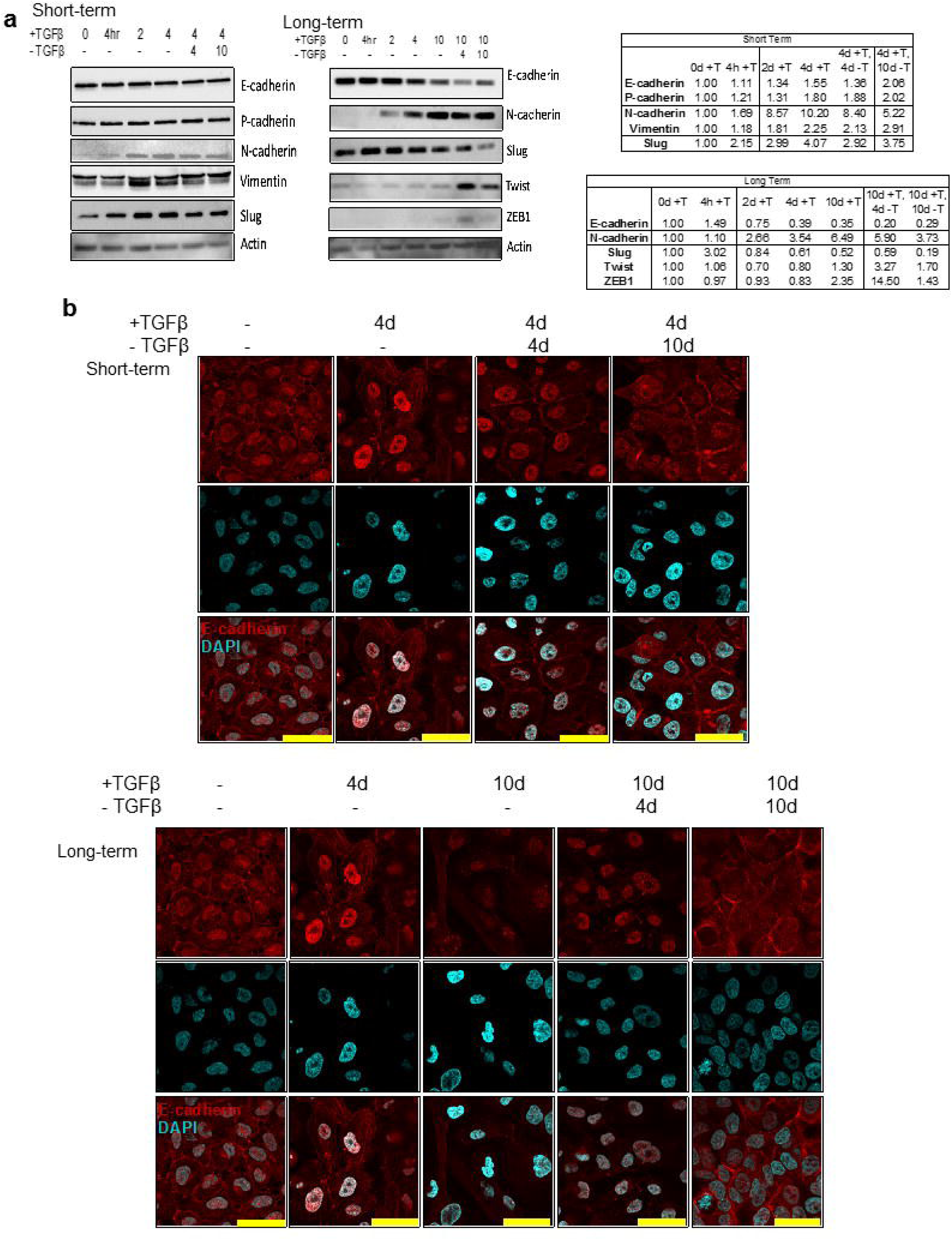
Protein expression dynamics during TGFβ treatment and withdrawal. **a** Western blot for epithelial and mesenchymal markers in MCF10A cells at indicated durations of 5 ng/mL TGFβ treatment and withdrawal in short-term (left) and long-term (right) TGFβ-induced EMT models. Table representing normalized values (relative to treatment loading and compared to untreated protein expression, respectively) shown to the right of graphs. **b** Representative confocal micrographs of E-cadherin localization (red) relative to nuclei (DAPI, blue) in short-term (top) and long-term (bottom) models. Scale bar = 30 µm.

Given the discordance between expression of *ZEB1* mRNA, upregulated at 2 days, and ZEB1 protein, detectable at 10 days, we next examined post-transcriptional regulation of *ZEB1* using a 3’UTR activity assay (23). MCF10A cells were transduced with a GFP-linked *ZEB1* 3’UTR reporter (GFP-Z1) and subjected to TGFβ treatment. Concomitant with the suppression of miR-200c and miR-203 (Fig. 1e), 2 days of TGFβ treatment was sufficient to observe elevated GFP-Z1 (Sup. Fig. 1a,b, green panel).

To better understand E-cadherin dynamics, we measured protein localization via immunofluorescence and FACS. Though the *CDH1*-RFP reporter indicated no changes in transcriptional activity in the short-term model (Sup. Fig. 1a,b, Tables 1,2), cell surface localization of E-cadherin was diminished at 4 days and remained low through 10 days of TGFβ treatment (Fig. 2b, Sup. Fig. 1a,b, purple panel). Strikingly, during TGFβ withdrawal, surface E-cadherin recovered in our short-term TGFβ treatment model, but not in our long-term TGFβ treatment model (Fig. 2b, Sup. Fig. 1a,b, purple panel). In the context of short-term TGFβ treatment, RNA-seq, protein expression, and the *CDH1-*RFP reporter, (Fig. 1d, 2a, Sup. Fig. 1a,b) do not indicate increasing E-cadherin expression during TGFβ withdrawal, while the immunofluorescence and FACS data (Fig. 2b, Sup. Fig. 1a,b) point to membrane localization. Such a recovery of surface E-cadherin suggests a cytoplasmic store capable of returning to the membrane following MET without the need for transcriptional upregulation. This finding is in concert with past studies on E-cadherin recycling (55).

In order to observe the relationship between TGFβ-induced EMT, TGFβ withdrawal-induced MET, and cellular plasticity, we characterized the mammosphere formation capacity of cells undergoing the long-term TGFβ treatment and withdrawal time course. Consistent with the observation that partial-EMT states exhibit maximal plasticity, seeding cells from such timepoints (4 days of treatment, or 10 days of treatment followed by 4 days of withdrawal) corresponded to significantly greater sphere formation than untreated cells or cells treated with TGFβ for 10 days (Sup. Fig. 2). Thus, inducing partial-EMT and partial MET stimulates cellular plasticity and non-adherent growth properties associated with formation of mammospheres.

### EMT and MET-induced chromatin accessibility regions

EMT is accompanied by reversible changes in the epigenome (19). Chromatin accessibility orchestrates dynamic gene expression by exposing or hiding regulatory genomic elements, resulting in distinct epigenomic states (56). To uncover the epigenomic basis associated with TGFβ-driven reversible EMT, we performed the assay for transposase-accessible chromatin with next-generation sequencing (ATAC-seq) (57). Given the diverse transcriptional dynamics observed in EMT-related genes, we first examined chromatin accessibility at these loci. Accessibility at *CDH1*, which remains high throughout TGFβ treatment but diminishes during withdrawal, was strikingly different from accessibility at *EPCAM*, which diminished as soon as 2 days of TGFβ (Fig. 3a, arrows). Notably, withdrawal in the short-term TGFβ treatment was characterized by a recovery of *EPCAM* accessibility that was not present following withdrawal following long-term TGFβ treatment (Fig. 3a, arrowhead). Additionally, *CDH3*, located immediately upstream of *CDH1*, follows similar accessibility patterns of *CDH1*, agreeing with observed protein expression patterns (Fig. 2a).

**Fig. 3.**
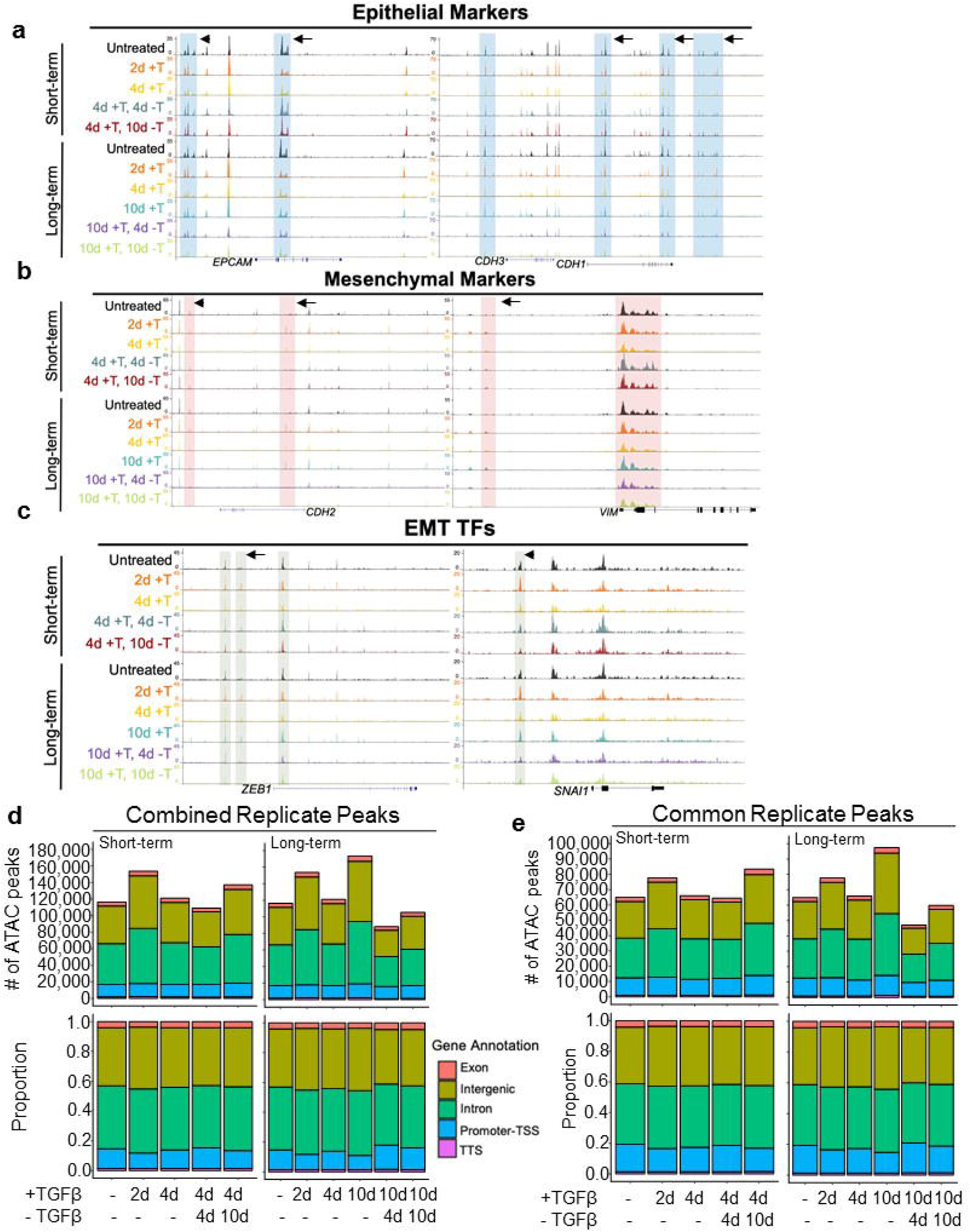
Dynamics of EMT- and MET-induced chromatin accessibility. ATAC-seq peak profiles of select: **a** epithelial (EPCAM, *CDH3*, and *CDH1*), **b** mesenchymal (*CDH2* and *VIM*), and **c** EMT-TF (*ZEB1* and *SNAI1*) genes in short-term (top) and long-term (bottom) TGFβ-induced EMT models (n = 2). Arrows and arrowheads indicate regions of interest. Number of accessible chromatin regions for **d** combined replicate peaks (top) and gene annotation proportions (bottom) in short-term (left) and long-term (right) models, and **e** number of peaks common among treatment replicates (top) and their gene annotation proportions (bottom) among replicates in short-term (left) and long-term (right) models (n = 2).

Accessibility at mesenchymal genes *CDH2* and *VIM* was more responsive to TGFβ treatment and withdrawal. In both models, treatment stimulated new accessibility regions in *CDH2* and upstream of *VIM* and these regions diminished following withdrawal (Fig. 3b, arrow). Interestingly, withdrawal following long-term TGFβ treatment also resulted in the dissolution of downstream ATAC-seq peaks previously prominent in untreated and TGFβ-treated conditions (Fig. 3b, arrowhead).

We also observed subtle chromatin accessibility alterations at EMT-TF genes *ZEB1* and *SNAI1*. New ATAC-seq peaks emerged upstream of the *ZEB1* gene following 2 days of TGFβ treatment (Fig. 3c, arrow). In both models, withdrawal diminished these peaks. *SNAI1* followed a similar pattern as upstream regions exhibited gains in accessibility following treatment which resolved, to a level lower than untreated cells, following withdrawal (Fig. 3c, arrowhead).

### EMT and MET proceed through distinct global epigenomic stages

We next examined the changes in chromatin accessibility from a global perspective by enumeration of all chromatin accessibility regions detected in the two replicates (Fig. 3d) and limiting the analysis to regions shared between the replicate treatments (Fig. 3e). The extent of chromatin accessibility was highly dynamic across both treatment models. Generally, TGFβ treatment was associated with greater chromatin accessibility while TGFβ withdrawal was associated with diminished accessibility (Fig. 3d,e). Dramatically, 10 days of treatment led to 50% more accessible sites than untreated cells (175,103 vs 113,680 comparing both replicate regions and 97,714 vs 65,238 in common replicate regions). This state was reversed upon 4 days of TGFβ withdrawal, whereupon nearly half of accessible sites were lost (91,335 sites comparing both replicate regions and 47,197 in common replicate regions) (Fig. 3d,e). Though the number of accessible chromatin regions fluctuated during reversible EMT, the regions remain similarly distributed amongst gene regulatory and intergenic loci (Fig. 3d,e). To stringently determine chromatin accessibility trends that accompany reversible EMT, we utilized regions common among both replicates for all subsequent analyses. To determine if TGFβ treatment resulted in the addition of new chromatin accessibility regions across the genome or near previously accessible sites, we interrogated the distance between ATAC-seq peaks and the number of ATAC-seq peak gaps exceeding 1 megabase (Mb). The median gap length was stable (medians 13,218-16,749 bp) throughout the timecourse with the exception of cells treated with TGFβ for 10 days (median 11,564 bp) and 10 days of TGFβ then withdrawal for 4 days (median 20,383 bp) (Fig. 4a). Concordantly, cells treated with TGFβ for 10 days exhibited nearly half the number of >1Mb gaps as other timepoints and cells undergoing TGFβ withdrawal had the most >1Mb gaps (Fig. 4b). Interestingly, when assessing the proximity of regions to known transcription start sites (TSSs), we observed that ATAC-seq peaks found in cells treated with TGFβ are further away from known TSSs (Sup. Fig. 3a, Table 3). These findings support the notion that newly accessible regions associated with 10 days of TGFβ treatment are being formed *de novo*, at locations distal to existing accessible regions, rather than clustered near to existing accessible regions.

**Fig. 4.**
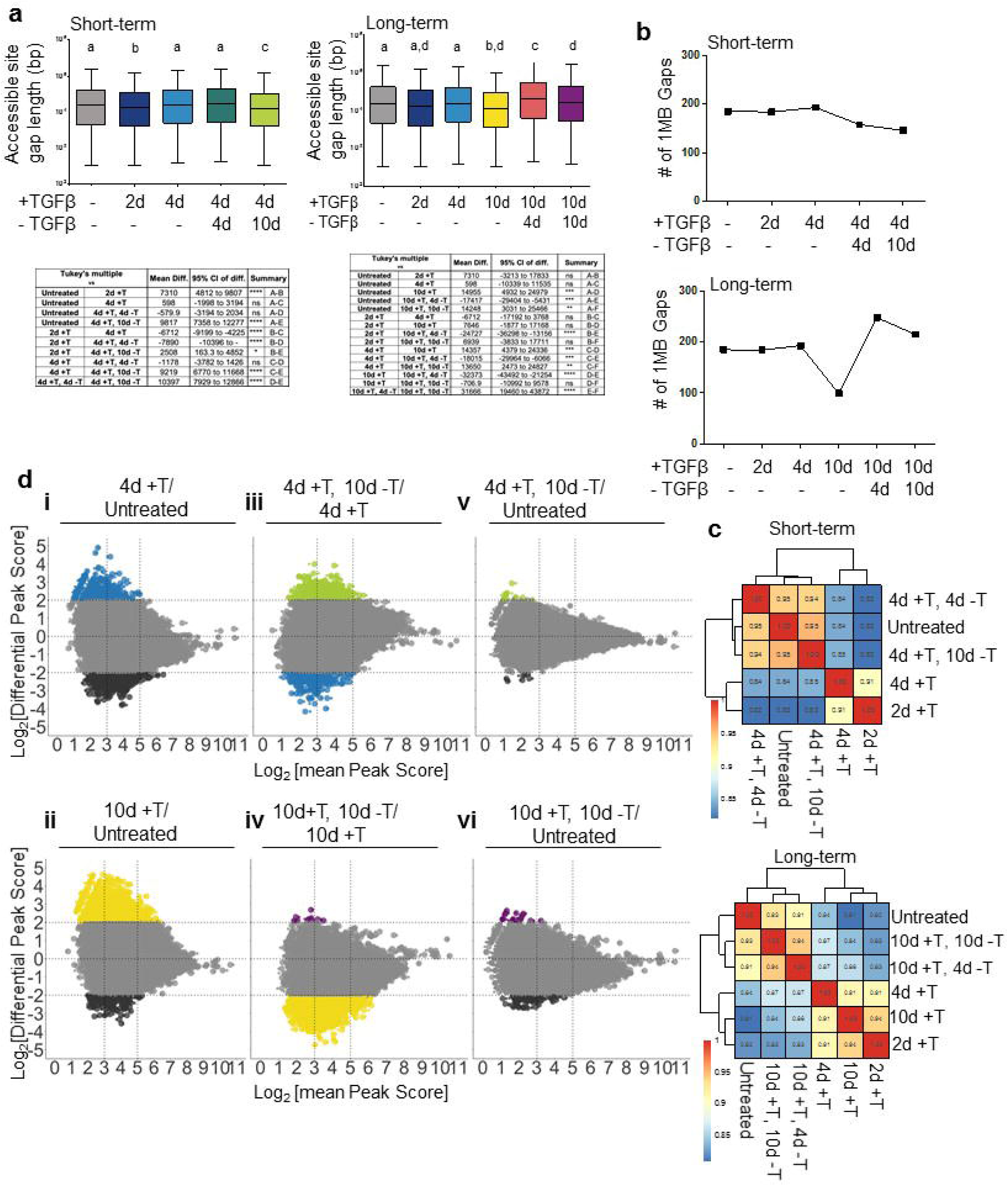
Differential accessibility is driven by changes at low- to moderately-accessible sites across the genome. **a** Gap length distribution (in bp) between ATAC peaks common among replicates in short-term (left) and long-term (right) TGFβ-induced EMT models. Data were analyzed using two-way ANOVA using Tukey’s multiple comparison test, statistical comparisons shown in table below graphs, ns = not significant, \**p* ≤ 0.05, \*\**p* ≤ 0.01, *** *p* ≤ 0.001, **** *<u>p</u>* < 0.0001. **b** Number of gaps between peaks common among replicates that exceed 1Mb in short-term (top) and long-term (bottom) models. **c** Pearson’s correlation analysis on quantile-normalized log-transformed ATAC peaks common among replicates in short-term (top) and long-term (bottom) models. **d** Differential accessibility (log_2_fold change in reads per 300-bp region) among selected EMT-induced and EMT-withdrawn conditions in short-term (top three) and long-term (bottom bottom) models. Colored peaks represent peaks at least 4-fold more accessibility at the indicated conditions. Dark gray = untreated, blue = 4d +TGFβ, light green = 4d +TGFβ, 10d -TGFβ, yellow = 10d +TGFβ, and purple = 10d +TGFβ, 10d -TGFβ.

Next, we assessed the similarities of chromatin accessibility profiles between timepoints using Pearson’s correlation coefficient of log_2_-transformed quantile-normalized peak scores. In both models, cells undergoing TGFβ treatment co-clustered while cells undergoing TGFβ withdrawal clustered together with untreated cells (Fig. 4c). The greatest distinctions in chromatin accessibility patterns were between the untreated and 2-day TGFβ treated cells (in short-term model, R = 0.82) and between the untreated and 10-day TGFβ treated cells (in long-term model, R = 0.81), suggesting that chromatin reprogramming occurs early, prior to many transcriptomic changes, and is sustained during EMT-induction. TGFβ withdrawal rapidly reverts DNA accessibility patterns near to untreated conditions (R = 0.94 and R = 0.95 in short- and long-term, respectively) (Fig. 4c), demonstrating that the epigenome is highly responsive to TGFβ treatment and withdrawal.

### Intensity and adjacent gene function of global epigenomic changes

We next considered whether changes in the chromatin accessibility regions were at peaks with low scores (log_2_ [mean Peak Score] <3), moderate scores (log_2_ [mean Peak Score] ≥3, <5), or high scores (log_2_ [mean Peak Score] ≥5), reasoning that, if timepoint-specific peaks were weak in intensity, the differences may be due to stochastic effects. When comparing untreated cells to cells exposed to TGFβ for 4 days or 10 days, peaks with the greatest difference in peak score (non-grey circles) were among the those with low to moderate peak score intensity values (Fig. 4d, i, ii). Additionally, while TGFβ results in an acquisition of new differentially-scored peaks, untreated conditions also contain some unique moderately-scored peaks which are lost following TGFβ treatment (Fig. 4d, i, ii, dark gray). Interestingly, short-term, rather than long-term, TGFβ treatment and withdrawal led to fewer differentially-scored peaks (Fig. 4d, iii, iv, light green and purple). When comparing untreated cells to cells that had undergone 10 days of TGFβ withdrawal from either model, ATAC-seq peaks with the greatest peak score difference (non-grey circles) were among the peaks with the overall weakest intensity values (Fig. 4d, v, vi; Sup Fig. 3b). These data indicate that the epigenomic dynamics associated with EMT and MET create and remove significant novel chromatin accessibility sites. Additionally, these data suggest that the grade of EMT induction following withdrawal may stimulate distinct MET chromatin states and epigenomes.

To better understand the phased implementation of EMT-associated chromatin accessibility, we enumerated the overlapping ATAC-seq peaks. As expected, many chromatin accessibility regions in untreated cells are present at other timepoints (n = 47,354) (Sup Fig. 4a). However, TGFβ treatment for any duration was associated with an additional 9,028 shared accessible regions, while a 10-day TGFβ treatment yielded an additional 16,164 unique regions (Sup Fig. 4a). Interestingly, cells which had been exposed to TGFβ, regardless of withdrawal time, retained some EMT-induced regions (Sup. Fig. 4b and 4c), despite the considerable chromatin constriction (Fig 3d,e). We then investigated the gene expression patterns of the subset of persistent EMT chromatin accessibility regions following long-term treatment and annotated the peaks to the closest gene and identified their gene expression patterns. Though chromatin accessibility was altered at these sites, most, but not all, gene expression returned to untreated conditions (Sup. Fig 4d, black box).

To derive the functional implications of these chromatin accessibility regions. we performed gene set enrichment analysis (GSEA) for the annotated genes closest to ATAC-seq peaks. msigDB hits revealed enrichment for Hallmark gene sets related to EMT, TNF-11 signaling, hypoxia, and TGFβ signaling as well as gene sets related to stemness and EMT in mammary cells or breast cancer (Sup. Fig. 5a, yellow). Enrichment of these gene sets diminishes following TGFβ withdrawal (Sup. Fig. 5a, purple). We were also interested in the function of genes near sites which remained accessible despite TGFβ withdrawal at our “Persistent EMT Peaks”. As above, this set was enriched for EMT, apical junction and mammary stemness genes(Sup. Fig. 5b). Notably, many of these genes remain expressed following 10 days of TGFβ withdrawal (Sup. Fig. 4d), constituting a potential signature of past EMT events.

### Dynamic transcription factor engagement during EMT and MET

We next hypothesized that TGFβ treatment would modulate enrichment of transcription factor binding sites (TFBSs) to enable partial or full EMT. We used HOMER motif discovery analysis to determine the enrichment of TFBSs found within ATAC-seq peaks (58). The degree of significance for motif enrichment was highly variable in both models (Fig. 5a). Consistent with the TGFβ-driven increase in the number of accessible sites (Fig. 3d), TGFβ treatment drove an increase in the overall significance of motif enrichment, which was diminished upon TGFβ withdrawal (Fig. 5a). Notably, the motifs which are highly significant in TGFβ-treated conditions were nonetheless enriched in other conditions (Sup. Fig 6a and b), indicating that introduction of novel TFBSs does not greatly contribute to the change in chromatin accessibility observed during reversible EMT.

**Fig. 5.**
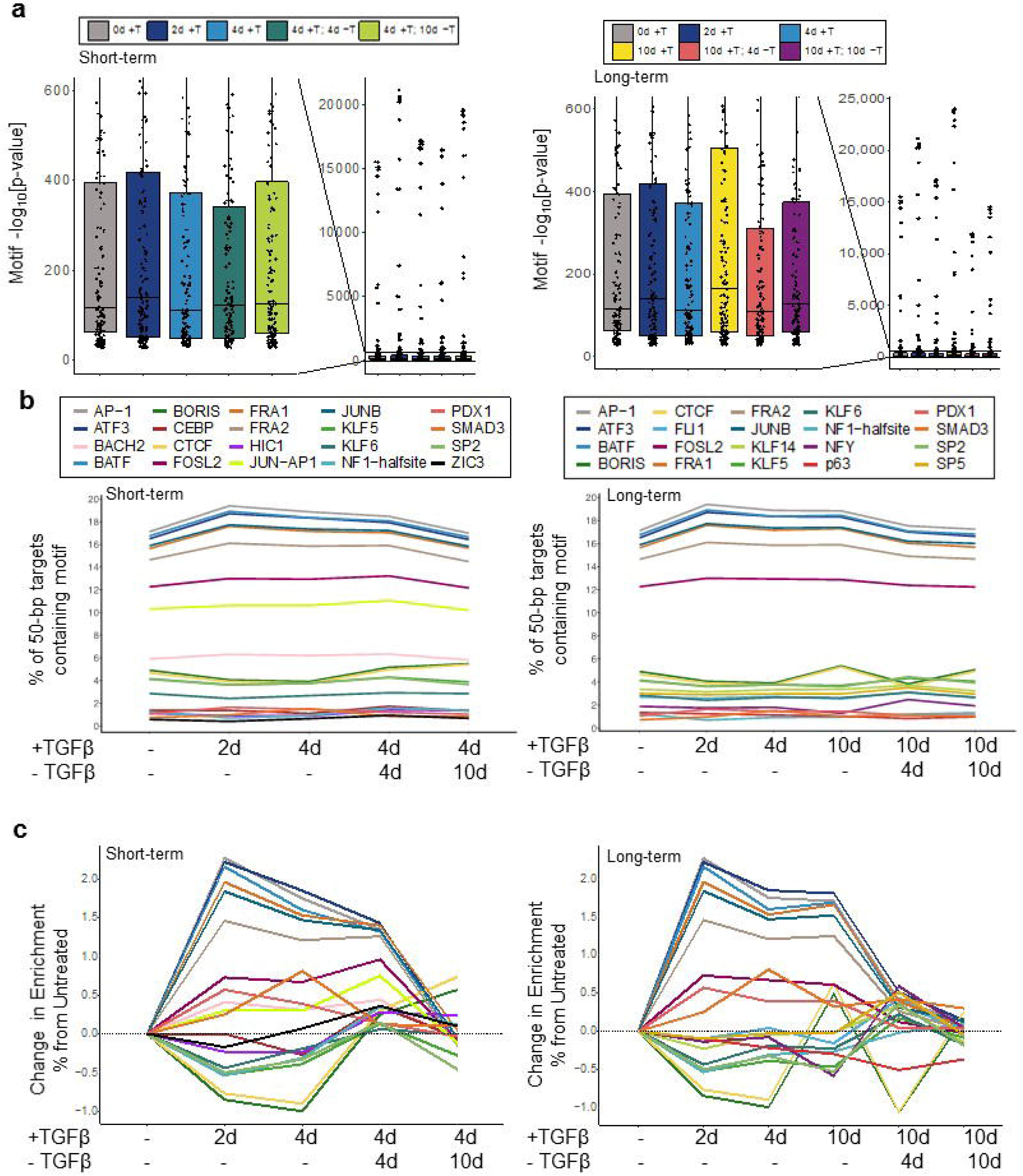
TFBS dynamics associated with increased accessibility and partial states. **a** Motif enrichment distribution for highly-significant (*p* ≤ 10^−12^) motifs in short-term (left) and long-term (right) TGFβ-induced EMT models. **b** Motif enrichment by percentage of 50-bp targets containing motif of the top 20 differentially-enriched motifs in short-term (left) and long-term (right) models. **c** Motif enrichment percent change from untreated of top 20 differentially-enriched motifs in short-term (left) and long-term (right) models.

We next wished to distinguish variation among individual transcription factors involved in progression through EMT and MET. We queried the percentage of 50-bp targets which contained the TFBS and limited our analyses to the top 20 enriched motifs. In both models, we noted considerable TGFβ-driven increases in the prevalence of AP-1 family and SMAD-binding motifs which resolved to baseline during TGFβ withdrawal (Fig. 5b). Because of the large range in TFBS prevalence, we calculated their percent change in comparison to untreated cells (Fig. 5c). While most TFBSs exhibited increased accessibility during TGFβ treatment, we observed a tri-modal pattern of enrichment for CTCF and BORIS motifs, with distinct enrichment in samples that were untreated, experienced 10 days of TGFβ, or experienced 10 days of TGFβ withdrawal (Fig. 5b,c).

### Loss of CTCF occupancy and protein accompanies partial-EMT and -MET states

CTCF and BORIS (*CTCFL*) have some overlapping functions and participate in establishing topologically associated domains, genetic imprinting, and gene expression (59, 60). However, as BORIS is not expressed in breast cancers and mammary cell lines (61), we further investigated the pattern of dynamically-engaged CTCF binding motifs. The distance between engaged CTCF binding motifs remained stable during the initial phases of TGFβ treatment but dropped significantly at 10 days of TGFβ treatment, rose upon 4 days of TGFβ withdrawal, and then stabilized to untreated levels at 10 days of withdrawal (Fig. 6a). As with the increase in general chromatin accessibility (Fig. 4a), the pattern of accessible regions containing CTCF motifs was consistent with novel engagement of CTCF bindings sites across the genome rather than in clusters.

**Fig. 6.**
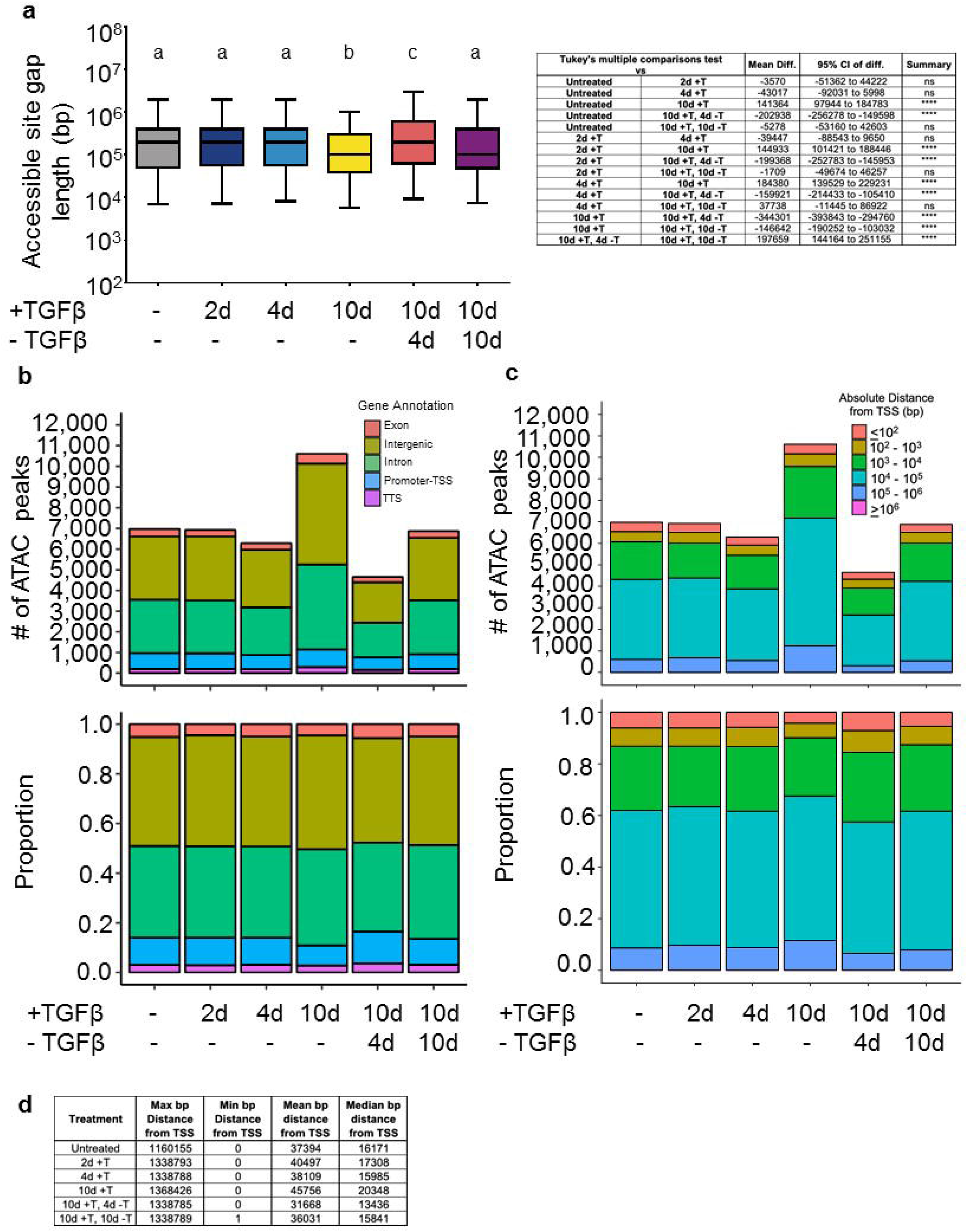
CTCF dynamic engagement is responsible for increased intergenic accessibility. **a** Gap length distribution (in bp) between ATAC peaks containing CTCF motifs (left). Table representing statistical comparisons (right). Data were analyzed using two-way ANOVA using Tukey’s multiple comparison test, statistical comparisons shown in table to the right of graph, **** *<u>p</u>* < 0.0001. **b** Table representing maximum, minimum, mean, and median bp distance from TSS for CTCF motif-containing accessible regions of peaks common among replicates in long-term model. **c** Number of accessible chromatin regions containing CTCF motifs (top) and their gene annotation proportion (bottom) in long-term model. **d** CTCF motif distance from TSS (in bp) for peaks containing CTCF motif (top) and their gene annotation proportion (bottom) in long-term model.

We next profiled the regulatory function of the target sites putatively occupied by CTCF. Ten days of TGFβ treatment was characterized by an increase in the proportion of CTCF binding motifs at intergenic loci and a decrease in the proportion at promoters (Fig. 6b, bottom panel), further supporting the notion that CTCF is binding to sites across the genome. Additionally, the majority of CTCF motifs were found within 20 kb of the nearest TSS and the most considerable changes for CTCF binding were at locations between 10 kb and 1 Mb from TSSs (Fig. 6c,d). Consistent with the role of CTCF as a regulator of long-distance chromatin interactions, these data suggest that, during EMT and MET, CTCF dynamically engages gene-distal sites with potential regulatory effects.

In order to understand the potential functional impacts of altered CTCF binding, we performed GSEA for the genes nearest to dynamically engaged CTCF binding sites. In long-term reversible EMT, most genes adjacent to CTCF motifs were shared amongst the three conditions in which CTCF engagement was enriched (n = 4,117) however 10 days of TGFβ treatment exhibited the largest number of characteristic genes (n = 1,518) (Sup. Fig. 7a). Interestingly, cells treated with TGFβ appear to retain some remnants of CTCF reconfiguration despite withdrawal (n=766) (Sup. Fig. 7a), highlighting a possible residual effect of EMT/MET. GSEA also reveals a connection between genes nearest to CTCF sites and cell development and chromatin biology, as bivalent (H3K4me3+H3K27me3) domains were highly enriched (Sup. Fig. 7b, Table 6).

Differential CTCF engagement throughout a timecourse of TGFβ treatment and withdrawal may be due to oscillations in CTCF expression. Despite considerable changes in motif enrichment, we note no changes in steady state *CTCF* mRNA levels (Fig. 7a). Conversely, western blotting revealed dynamic protein expression. In untreated conditions, Upon four days of TGFβ treatment, CTCF expression decreases until around 8 days of withdrawal (Fig. 7b). Given the link between CTCF and both bivalent domains (Sup. Fig. 7b, Table 6) and LOCK domains, we assessed the global levels of relevant histone modifications. Concomitant with CTCF repression patterns, we also observed increases in H3K4-, H3K27-, and H3K9-trimethylation marks (Fig. 7b). We hypothesized that TGFβ treatment could induce a proteasome-mediated degradation of CTCF, however treatment with TGFβ and proteasome inhibitors did not rescue CTCF expression (data not shown). Collectively these data demonstrate that CTCF is dynamically regulated at the protein level during partial and reversible EMT and suggests that CTCF protein levels largely dictate the accessibility of CTCF binding motifs.

**Fig. 7.**
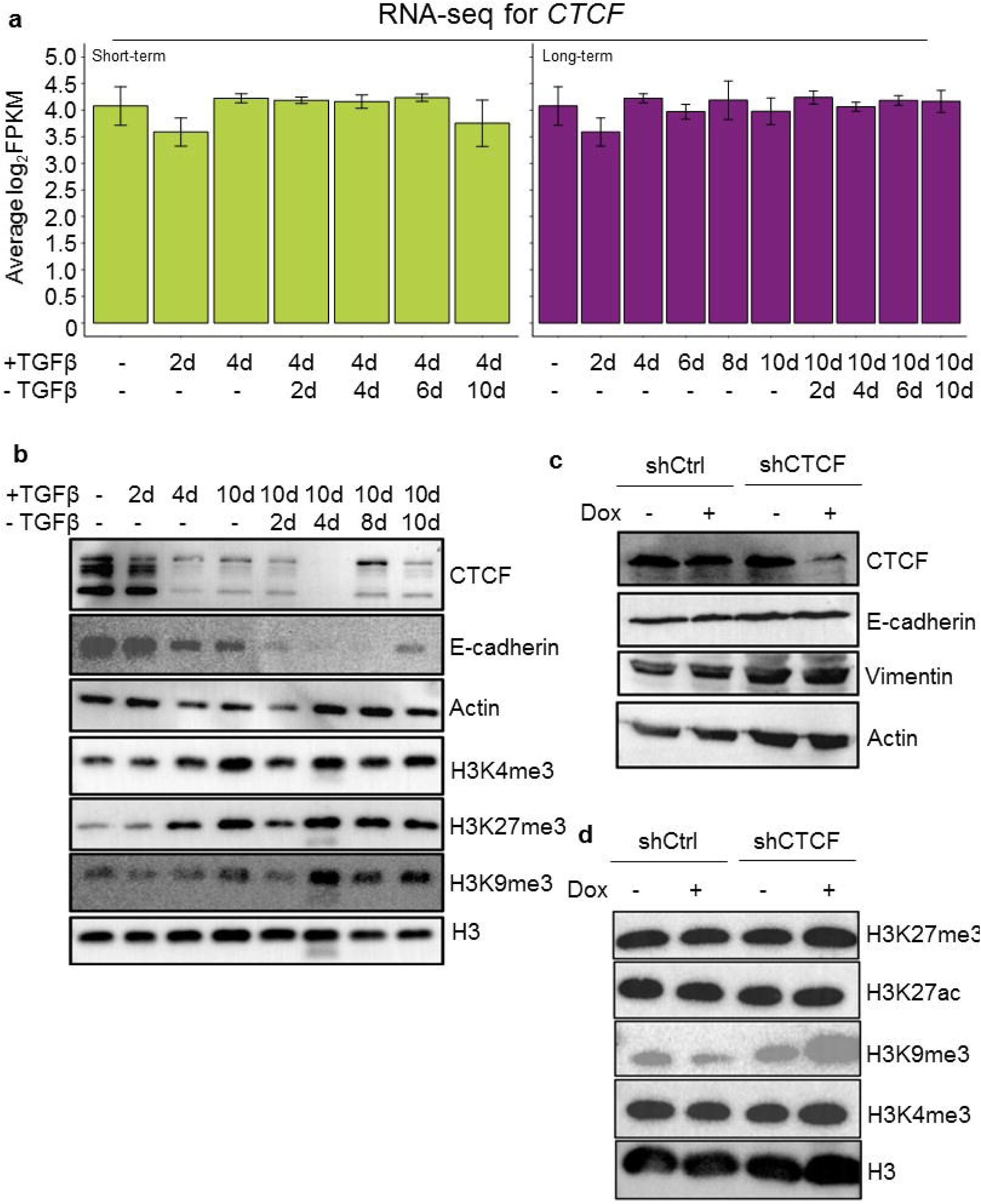
CTCF protein and histone modifications are dynamically expressed during reversible EMT and CTCF repression is accompanied by upregulation of H3K9me3 marks. **a** Average log_2_FPKM counts for *CTCF* gene based on RNA-seq in short-term (left, lime green) and long-term (right, purple) TGFβ-induced EMT model (n = 2, mean ± SEM). **b** Western blot for CTCF, E-cadherin, and various H3 post translational modifications in long-term reversible EMT model. **c** Western blot for protein expression for epithelial and mesenchymal markers and histone tail modification expression following 3d 0.5 µg/mL doxycycline on inducible short hairpin CTCF knockdown or non-targeting control in MCF10A cells. **d** Western blot for protein expression for epithelial and mesenchymal markers and indicated histone H3 post-translational modifications following 4 days of 0.5 µg/mL doxycycline on inducible short hairpin CTCF knockdown or non-targeting control in MCF10A cells.

We sought to explore the implications of CTCF repression in EMT and MET induction. To do so, we generated doxycycline-inducible shRNA-expressing MCF10A cells to determine the effect of CTCF repression on EMT marker and histone modification levels. Surprisingly, CTCF repression alone was not sufficient to induce EMT in MCF10A cells, failing to alter the expression of E-cadherin or vimentin (Fig. 7c). However, CTCF repression was sufficient to increase global levels of H3K4me3 and H3K27me3 (Fig. 7d), corroborating the association between CTCF motifs and bivalent domains (Sup. Fig. 7b). Additionally, we noted increased H3K9me3 levels associated with TGFβ-induced CTCF repression (Fig 7d), pointing to a potential effect on H3K9me3-mediated gene silencing in LOCK domains. Overall, these data demonstrate that CTCF protein, not mRNA, is dynamically regulated during long-term reversible EMT and that loss of CTCF drives alterations to the histone code.

## Discussion

EMT and its reversal, MET, are important for normal physiological processes such as development and wound healing and is hypothesized to endow cancer cells with metastatic abilities. EMT has been revealed to progress through partial states in terms of morphology, gene expression, and expression of epithelial protein marker E-cadherin. In studies utilizing solid tumor models, it has been shown that cells within an intermediate-mesenchymal state are more spheroidogenic and resistant to anoikis (43) and cells that express both KRT14 and vimentin disproportionately contribute to metastasis (40). Despite this importance, the factors mediating the progression through partial-EMT states and back are not well characterized. In this study, we have extensively characterized the chromatin accessibility alterations and gene expression output that occur during distinct stages of reversible TGFβ-induced EMT.

In our study, we demonstrate that short-term TGFβ elicits dynamic trafficking of E-cadherin to and from the cell membrane. Four days of TGFβ treatment reduces the proportion of cells with detectable surface E-cadherin despite a minor repression of overall E-cadherin protein or transcript. This finding corroborates a report by Aiello et al. who observed that pancreatic cancer cells with partial-EMT features exhibit high levels of intracellular E-cadherin which co-localizes with recycling endocytic vesicles (54). However upon 10 days of TGFβ treatment we observed the emergence of cells which have not only lost membranous E-cadherin but show a diminishment in their total E-cadherin protein and *CDH1* accessibility. This finding was unexpected and suggests that persistent exposure to EMT-inducing stimuli imposes chromatin alterations that are not reversed despite withdrawal of the stimulus. While outside the bounds of the present study, EMT-induced DNA methylation has been implicated in stable mesenchymal-like phenotypes by repression of miR-200b, miR-200c, miR-203 and other targets (20, 21, 23, 30, 62).

Transcriptomic determination of EMT status utilizing an EMT-specific gene expression signature has linked EMT enrichment to claudin-low breast cancers (34), mesenchymal glioblastoma (63), rapamycin-resistance in breast cancer (64), radio-resistance in prostate cancer (65), and response to immune checkpoint blockade therapy in lung cancer (66). (35, 44, 47, 54, 67). Herein, we show that transition through both EMT and MET involves stage-wise changes in EMT-signature gene expression. Moreover, TGFβ-withdrawal from cells treated for only 4 days elicited changes in gene expression that largely mirrored patterns observed at 4 days of treatment. However, withdrawal following 10 days of TGFβ treatment led to prolonged expression of N-cadherin and Twist and failure to fully re-express E-cadherin. These data are similar to those recently reported by Jia et al. who find that the ZEB1 3’UTR reporter remains expressed despite prolonged TGFβ exposure and withdrawal (68). It has been also reported that partial-EMT states are coordinated by Slug (*SNAI2*) while Snail (*SNAI1*) and ZEB-1 activation can promote a mesenchymal phenotype (69). Our results agree with this observation, as Slug protein expression increases as early as 4 hours of TGFβ exposure while ZEB1 protein expression is observed only following 10 days of TGFβ and 4 days of withdrawal, at the time point which expresses the least amount of E-cadherin, *CDH1*, and *CDH1-*RFP activity.

We show that EMT is accompanied by dynamic increases in chromatin accessibility. These observations are in-line with prior observations of murine mammary epithelial cells in response to TGFβ treatment (70). We note that striking chromatin alterations occur relatively rapidly following 2 days of TGFβ as 22,354 nascent peaks appear while only 9,678 peaks are lost.

Many of the newly-acquired regions have low accessibility scores and may contribute to some of the stochastic transcriptional noise that permits cell plasticity(71). Overall TGFβ withdrawal-induced MET causes a decrease in accessibility inducing a global chromatin “shutdown” whereupon the number of accessible regions is reduced by half and the distance between remaining regions significantly increases.

We identify that several factors increase their degree of chromatin engagement including SMAD and AP-1 transcription factors, which have been previously reported to mediate and be regulated by EMT (39, 40, 48, 70, 72). We interpret these results to suggest that chromatin is highly responsive to TGFβ treatment, permitted through numerous relatively weak interactions between a broad set of DNA-binding proteins that create a permissive chromatin environment. Following TGFβ withdrawal, these DNA-binding proteins lack the signals to sustain their occupancy, and the chromatin compacts until the cell re-establishes epigenetic homeostasis similar to baseline conditions, Though most transcription factors were responsive to TGFβ treatment or withdrawal, interestingly we observed tri-modal enrichment of CTCF motifs and declines during transitional, or partial-EMT and -MET states. CTCF participates in gene expression regulation, chromatin organization and genetic imprinting. In cancer, CTCF-binding sites have been found to be mutated in solid tumors (59, 60) and methylation at CTCF binding elements has been shown to drive overexpression of oncogenes such *PDGFRA*(73). This dynamic enrichment is potentially driven by fluctuations in CTCF protein expression or stability. Notably, Pastushenko et al., which describes extant populations of cells in partial EMT states, applied ATAC-seq to highly characterized, sorted populations of epithelial (Epcam+), mesenchymal (CD106^+^/CD51^+^/CD61^+^), and intermediate (CD51^-^/CD61^-^) SCC cells (40). Motif enrichment analysis of ATAC-seq data from Pastushenko et al. also demonstrates enrichment for CTCF binding motifs in accessible sites in epithelial and highly-mesenchymal states and a reduction in enrichment in partial-EMT states (40). Additionally CTCF has been shown in ovarian cancer to promote cancer metastasis by increasing the expression of genes such as *CTBP1, SERPINE1*, and *SRC* (74); however we did not observe accessible CTCF motifs adjacent to these genes in our model. A recent report by Li et al. demonstrates that an alternatively-spliced variant of CTCF can occupy canonical CTCF sites and cause differential effects on chromatin architecture gene expression (75). Therefore, it would be interesting to distinguish if alternatively-spliced variants of CTCF participate in partial stages and account for divergent effects observed between our model and others.

## Conclusions

In conclusion, our study characterizes 1) E-cadherin expression and localization dynamics, 2) partial-EMT/MET gene expression, 3) chromatin alterations occurring during reversible EMT, 4) expression patterns and motif enrichment of CTCF, and 5) the link between CTCF repression and histone modifications. We conclude that E-cadherin dynamics are distinct in partial-EMT and partial-MET, and that CTCF loss facilitates chromatin reorganization in partial-EMT/MET states while re-expression drives the establishment of novel chromatin arrangements leading to altered gene regulation in both EMT and MET conditions, linking CTCF dynamics to epithelial-mesenchymal plasticity.

## Methods

### Cell culture

MCF10A cells (ATCC) were cultured in DME/F12 media (GE Healthcare Life Sciences) supplemented 5% horse serum (GE Healthcare Life Sciences), 1% penicillin/streptomycin (Lonza), 20 ng/mL EGF (Sigma), 10 ng/mL insulin (Sigma), 500 ng/mL hydrocortisone (Acros Organics), and 100 ng/mL cholera toxin (Enzo Life Sciences). Cells were plated at 10,000 cells/cm^2^ and passaged every other day to maintain consistent densities. For TGFβ treatment, media was supplemented with 5 ng/mL recombinant human TGFβ-1 (R&D Systems; resuspended in 4 mM HCl, 0.1% BSA).

### Viral transductions

Stable Z-cad reporter cell lines were generated via viral transfection as described (23). In short, MCF10A cells were dually transduced with pHAGE-E-cadherin-RFP and FUGW-d2GFP-ZEB1 3’ UTR lentiviruses packaged by transfected HEK-293T cells. RFP^+^/GFP^+^ doubly labeled MCF10A cells were sorted via FACSMelody™ Cell Sorter (BD Biosciences).

For viral transfection of doxycycline-inducible shCTCF and shCtrl cell lines, HEK-293T cells (ATCC) were cultured in DMEM (Corning) supplemented with 10% FBS (Equitech-Bio) and 1% penicillin/streptomycin (Lonza). Virus DNA liposomes were prepared using FuGENE HD Transfection reagent (Promega) and DMEM. Vector-expressing cells were selected with 10 μg/mL puromycin.

### Western blotting

Cells were harvested and resuspended in RIPA buffer (Alfa Aesar) supplemented with protease and phosphatase inhibitors (ThermoScientific) and incubated in ice for 60 minutes. Lysed cells were centrifuged at 15,000 rcf for 20 minutes at 4° C and the supernatant was isolated. Protein concentrations were determined using a bicinchoninic acid assay (ThermoScientific). Approximately 40 μg of protein per treatment was loaded into a 12% SDS-polyacrylamide gel and run at 125 V for 90 minutes. Following electrophoresis, gels were transferred to methanol-activated PVDF membranes (Thermo Scientific) for 2.5 hours at 0.250 A, and blots were blocked in 5% milk (Carnation) in TBST for 30 minutes at room temperature. Primary antibodies were incubated overnight at 4°C and secondary antibodies were incubated at room temperature for 1 hour. Chemiluminescence signal was obtained using ECL™ Prime (GE Healthcare) and blot images were generated using the ChemiDoc™ Imaging System (Bio-Rad).

### Flow cytometry

At the conclusion of the timecourse, 2.5 x 10^6^ Z-CAD reporter cells were counted and resuspended in 500 μL 1% FBS (Equitech-Bio) in PBS with anti-E-cadherin BV421 (#745965, 1:100, BD Biosciences) and incubated on ice for 90 minutes. Following incubation, cells were pelleted and washed twice with 1% FBS in PBS and subjected to flow cytometry using a FACSMelody™ (BD Biosciences).

### Immunostaining

Live cells plated on glass coverslips were fixed with 2% paraformaldehyde (Acros Organic) in PBS (Lonza) for 20 minutes at room temperature, followed by washing with PBS. Fixed tissues were permeabilized with 0.3% Triton X-100 (Fisher Bioreagents) in PBS for 15 minutes followed by a series of PBS washes. Cells were quenched with 1% glycine (Fisher Bioreagents) in PBS, washed with PBS, and blocked in 8% BSA (Alfa Aesar) in TBST at 4°C overnight. Coverslips were incubated in primary antibody at 4°C overnight. Following incubation, coverslips were washed in TBST and subjected to secondary antibody dilution for 1 hour at room temperature. Nuclei were stained with DAPI for 5 minutes, washed with DNase and RNase-free water, then affixed to glass slides with Permount (Fisher Scientific).

Brightfield imaging was performed using a Nikon Eclipse Ts2R microscope with DS-Qi2 camera and analyzed, scaled, and edited using Nikon NIS Elements v4.5 Imaging Software. Confocal immunofluorescence imaging was performed using an Olympus FV-3000 Confocal Laser Scanning Microscope and analyzed, scaled, and edited using Olympus FV31S-SW software. Image cropping was performed using ImageJ (version 1.51s).

### Antibodies

The following antibodies and dilutions (in 5% milk in TBST) were used for western blotting: E-cadherin (#14472, monoclonal mouse, 1:1000, Cell Signaling), N-cadherin (#13116, monoclonal rabbit, 1:1000, Cell Signaling), P-cadherin (#2189, monoclonal rabbit, 1:1000, Cell Signaling), Vimentin (#10366-1-AP, polyclonal rabbit, 1:2000, ProteinTech), Slug (#9585, monoclonal rabbit, 1:1000, Cell Signaling), CTCF (#07-729, polyclonal rabbit, 1:1000, Millipore Sigma), Actin (#612656, monoclonal mouse, 1:2000, BD Biosciences), rabbit HRP-linked IgG secondary (#7074S, 1:2000 Cell Signaling), mouse HRP-linked IgG secondary (#926-80010, 1:2000, Li-Cor).

The following antibodies and dilutions (in TBST for primary or 3% BSA in TBST for secondary) were used for immunofluorescence: E-cadherin (#14472, monoclonal mouse, 1:100, Cell Signaling), mouse-IgG Alexa Fluor 594-conjugated secondary (#8890, 1:100,000, Cell Signaling).

### RNA extraction and qPCR

RNA was extracted from cell cultures using TriZol Reagent (Thermo Fisher) and isolated following manufacturer guidelines. 500 ng of RNA was used for input for cDNA synthesis using miR-203, miR-200c, and U6-specific TaqMan Gene Expression primers (Applied Biosystems). Quantitative PCR analyses were performed using cDNA as a template, TaqMan-specific qPCR primers, and TaqMan Gene Expression Master Mix (Applied Biosystems). QuantStudio5 real-time PCR machine (Applied Biosystems) with four technical replicates per biological replicate. Signal was quantified and normalized using QuantStudios5 software (version 1.5.1).

### Mammosphere Assays

At the conclusion of the timecourse, 5 × 10^3^ cells were plated in low-attachment 96-well plates containing 100 μL MEGM media (without BPE) (Lonza, CC-3150), 20 ng/mL bFGF (Sigma Aldrich), 10 ng/mL EGF (Sigma Aldrich), 4 μg/mL heparin (Sigma Aldrich), and 1% methylcellulose. 25 μL mammosphere media was added to each well every third day. Spheres were allowed for form for 14 days.

### RNA-seq Library Preparation and Sequencing

RNA was extracted from cell cultures using TriZol Reagent (Thermo Fisher) and isolated following manufacturer guidelines. Libraries were prepared using TruSeq Stranded mRNA Library Prep Kit (Illumina).

### EMT score calculation

The EMT scores were calculated utilizing the 76-gene expression signature reported (51) and the metric mentioned based on that gene signature (76). For each sample, the score was calculated as a weighted sum of 76 gene expression levels, and weights were measured based on the correlation of a particular gene with CDH1 expression. The scores were standardized for all the samples in the dataset by subtracting the mean across samples so that the global mean of the score was zero. Negative scores calculated using this method can be interpreted as mesenchymal phenotype and the positive scores as epithelial.

### ATAC-seq Library Preparation and Sequencing

ATAC-seq libraries were generated as previously described (57). Briefly stated, 5 × 10^4^ cells obtained from cell culture, centrifuged and cell pellets were resuspended in 50 μL lysis buffer μ(10 mM Tris, 10 mM NaCl, 3 mM MgCl_2_, and 0.1% IGEPAL CA-630) and centrifuged at 500g for 10 min at 4°C. The pellet was resuspended in transposase reaction mix (25 uL 2X TD buffer, 2.5 μL Transposase (Nextera DNA sample preparation kit, Illumina), and 22.5 μL H_2_0 and incubated at 37°C for 30 min. Tagmented DNA was purified using MinElute PCR Purification Kit (Qiagen) per manufacturer’s instructions. DNA libraries were PCR-amplified using Nextera DNA Sample Preparation Kit (Illumina) using the following PCR conditions: 98°C for 30s, then thermocycling for 98°C for 10s, 63°C for 30s, and 72°C for 1 min for 12 cycles followed by 72°C for 5 min. PCR products were size-selected for 200 to 800 base pair fragments using SPRI-Select Beads (Beckmann-Coulter). ATAC-seq reads were paired-end sequenced using an Illumina NextSeq500.

### ATAC analysis

Due to the similarities of both ends, only one paired-end read was used for analysis. Adapter sequences were removed using cutadapt (Galaxy version 1.16.6) and reads were cropped to 30 bp with Trimmomatic (Galaxy Version 0.36.6). ATAC-seq reads were aligned to *hg19* using Bowtie2 (v2.3.4.2). Mitochondrial, unmapped and random contigs, and ChrY reads were excluded using samtools (v1.9) (77) to generate filtered bam files.

Tag directories were generated in HOMER from filtered bam files (58). Peaks from each replicate were called separately using HOMER by calling ‘findPeaks [Tag Directory] -style dnase -o auto.’ Following, replicate peaks were merged by calling ‘mergePeaks -d 300 [Rep 1] [Rep 2]’ to determine common peaks. Common peaks were used for downstream analyses.

For Pearson correlation analyses, combined tag directories containing both replicates were generated to quantify average peak score. Common merged peaks from each replicate were merged with other time course conditions by calling ‘mergedPeak -d 300 [Peak files].’ Time course peak files were annotated and quantified to hg19 by calling “annotatePeaks.pl [Merged Peak file] hg19 -size 300 -log -d.” Peak scores for each condition were quantile-normalized by the preprocessCore package (v1.44.0) in R (v3.5.1). Quantile-normalized Pearson correlation scores were generated by R Base and visualized by the pheatmap R package (v1.0.12).

*De novo* and known motif searches were performed using HOMER (Heinz et al., 2010) for 50 basepair regions excluding masked genomic regions by calling ‘findMotifsGenome.pl [Common peaks] hg19 -size 50 -mask.” Motifs with p-values > 10^−12^ were discarded. Plots representing the number of significant motifs (p-value ≤ 10^−12^) and their enrichment were generated using the ggplot2 R package (3.2.0).

### Venn Diagrams and GSEA analysis

Peak similarities were identified using mergePeaks in HOMER and annotated to *hg19* to produce the Entrez IDs for the gene promoters nearest to peaks. Entrez IDs were input in Venn Diagram tool (http://bioinformatics.psb.ugent.be/webtools/Venn/) to enumerate similarities. Accessed 11 November 2019. LucidChart was used to produce Venn Diagrams. GSEA analysis for MSigDB hits was performed for specific peak groups. Because of the abundance of MSigDB hits, our analysis was limited to either unbiased Hallmarks-only gene sets (78), or breast and mammary-specific gene sets (containing keywords “breast” or “mammary”).

### Primers

The following primers were used for ATAC sequencing:

N501: AATGATACGGCGACCACCGAGATCTACACTAGATCGCTCGTCGGCAGCGTC

N502: AATGATACGGCGACCACCGAGATCTACACCTCTCTATTCGTCGGCAGCGTC

N701: CAAGCAGAAGACGGCATACGAGATTCGCCTTAGTCTCGTGGGCTCGGAGATGT

N702: CAAGCAGAAGACGGCATACGAGATCTAGTACGGTCTCGTGGGCTCGGAGATGT

N703: CAAGCAGAAGACGGCATACGAGATTTCTGCCTGTCTCGTGGGCTCGGAGATGT

N704: CAAGCAGAAGACGGCATACGAGATGCTCAGGAGTCTCGTGGGCTCGGAGATGT

N707: CAAGCAGAAGACGGCATACGAGATGTAGAGAGGTCTCGTGGGCTCGGAGATGT

N708: CAAGCAGAAGACGGCATACGAGATCCTCTCTGGTCTCGTGGGCTCGGAGATGT

N710: CAAGCAGAAGACGGCATACGAGATCAGCCTCGGTCTCGTGGGCTCGGAGATGT

N711: CAAGCAGAAGACGGCATACGAGATTGCCTCTTGTCTCGTGGGCTCGGAGATGT

N712: CAAGCAGAAGACGGCATACGAGATTCCTCTACGTCTCGTGGGCTCGGAGATGT

### Quantification and Statistical analysis

Two-tailed Student’s *t*-test and ANOVA were performed using GraphPad Prism (v6) and R (v 3.5.1).

### Availability of data and materials

The ATAC and RNA sequencing data generated during the current study are available in the NCBI Gene Expression Omnibus (GEO) and are accessible though GEO series accession number GSE145851 (https://www.ncbi.nlm.nih.gov/geo/query/acc.cgi?acc=GSE145851).

## Declarations

### Ethics approval and consent to participate

Not applicable

### Consent for publication

Not applicable

### Author Contributions

J.H.T. and Y.L. conceived, managed, and arranged funding for the project; J.H.T. and K.S.J. designed the experiments; K.S.J. performed most of the experiments and performed ATAC- and RNA-seq bioinformatics analysis; S.H., P.C., and M.K.J. provided computational analysis for RNA-seq; S.S. assisted with quantitative PCR data; K.S.J. and M.J.T. contributed to the Z-CAD transduced cells used in this study; Y.L. provided training and guidance with bioinformatic analyses. J.H.T. and K.S.J. wrote the manuscript. All authors read and approved the final draft of the manuscript.

## Acknowledgements

We thank Dr. Dwayne Simmons for generously permitting the use of his lab materials. We thank Dr. Michelle Nemec and the Baylor University Molecular Biosciences Center for support during the course of this work. We would like to thank Dr. Bernd Zechmann (Center for Microscopy and Imaging, Baylor University, Waco, TX) for technical support during microscopy and image analysis.

