## Supplement for "Gene expression and chromatin accessibility during progressive EMT and MET linked to dynamic CTCF engagement"

**a** Short-term

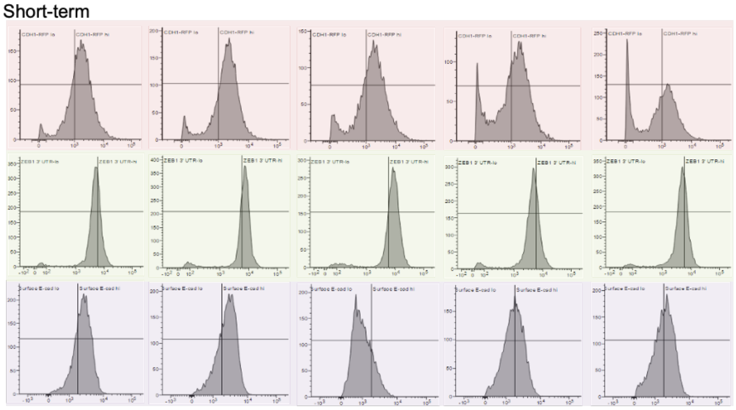

| | | | | | |
|---|---|---|---|---|---|
| +TGFβ | - | 2d | 4d | 4d | 4d |
| - TGFβ | - | - | - | 4d | 10d |

**b** Long-term

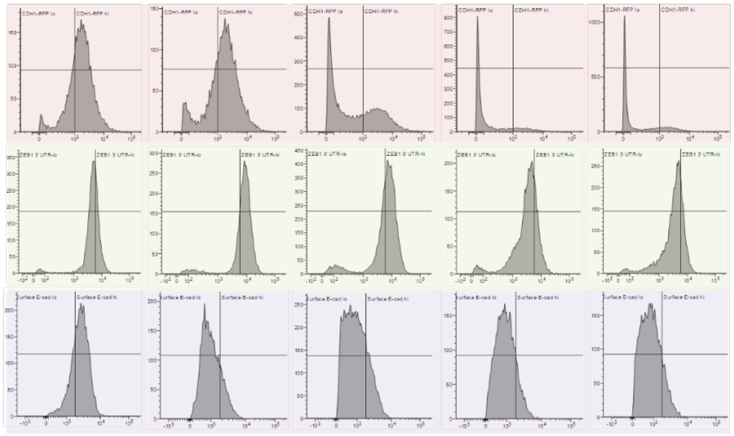

| | | | | | |
|---|---|---|---|---|---|
| +TGFβ | - | 4d | 10d | 10d | 10d |
| - TGFβ | - | - | - | 4d | 10d |

**Table 1** Short-term

| | Percent of Parent Population | | | | |
|---|---|---|---|---|---|
| | Un-treated | 2d +T | 4d +T | 4d +T, 4d -T | 4d +T, 10d -T |
| CDH1 Reporter hi | 72.4% | 77.6% | 69.6% | 58.7% | 49.8% |
| CDH1 Reporter lo | 27.6% | 22.4% | 30.4% | 41.3% | 50.2% |
| ZEB1 3' UTR Reporter hi | 31.2% | 67.1% | 77.4% | 27.4% | 30.5% |
| ZEB1 3' UTR Reporter lo | 68.8% | 33.0% | 22.6% | 72.6% | 69.5% |
| Surface E-cad hi | 72.0% | 71.3% | 18.6% | 45.6% | 51.8% |
| Surface E-cad lo | 28.0% | 28.7% | 81.4% | 54.4% | 48.3% |

**Table 2** Long-term

| | Percent of Parent Population | | | | |
|---|---|---|---|---|---|
| | Un-treated | 4d +T | 10d +T | 10d +T, 4d -T | 10d +T, 10d -T |
| CDH1 Reporter hi | 72.4% | 69.6% | 37.5% | 13.1% | 17.6% |
| CDH1 Reporter lo | 27.6% | 30.4% | 62.5% | 86.9% | 82.4% |
| ZEB1 3' UTR Reporter hi | 31.2% | 77.4% | 61.5% | 21.5% | 21.5% |
| ZEB1 3' UTR Reporter lo | 68.8% | 22.6% | 38.5% | 78.5% | 78.5% |
| Surface E-cad hi | 72.0% | 18.6% | 20.3% | 14.1% | 13.8% |
| Surface E-cad lo | 28.0% | 81.4% | 79.7% | 85.9% | 86.2% |

**Supplementary Figure 1.** FACS profiling of *ZEB1* 3' UTR reporter, *CDH1* promoter reporter, and surface E-cadherin in MCF10A cells. **a** FACS quantification for Z1-GFP reporter (green panel), *CDH1*-RFP reporter (pink panel), and surface E-cadherin (purple panel) in short-term TGFβ model (n = 1). **b** FACS quantification for Z1-GFP reporter (green panel), *CDH1*-RFP reporter (pink panel), and surface E-cadherin (purple panel) in long-term TGFβ model (n = 1). **Table 1** Percentage values for populations defined in Supp. Fig. 1a**. Table 2** Percentage values for populations defined in **b**.

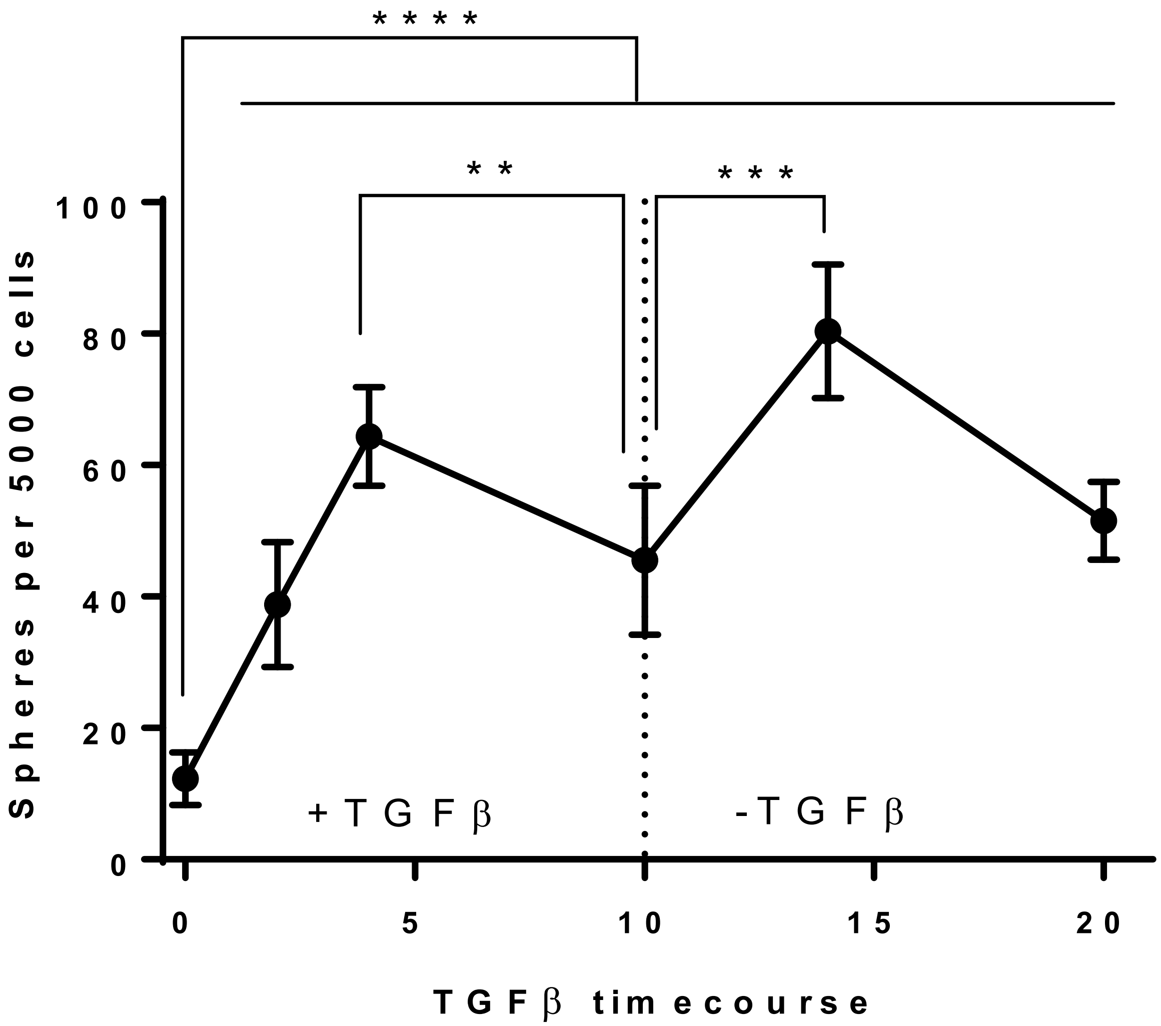


**Supplementary Figure 2.** Mammosphere-formation capacity of cells treated with TGFβ. Following the conclusion of the time course, TGFβ-treated and -withdrawn cells were subjected to mammosphere-promoting conditions and mammospheres $\geq$ 50 μm were counted after 10 days. Data were shown as mean + SEM from n = 5 with the indicated significance by using a two-tailed Student's *t* test, **$p < 0.01$, *** $p < 0.001$, **** $p < 0.0001$.

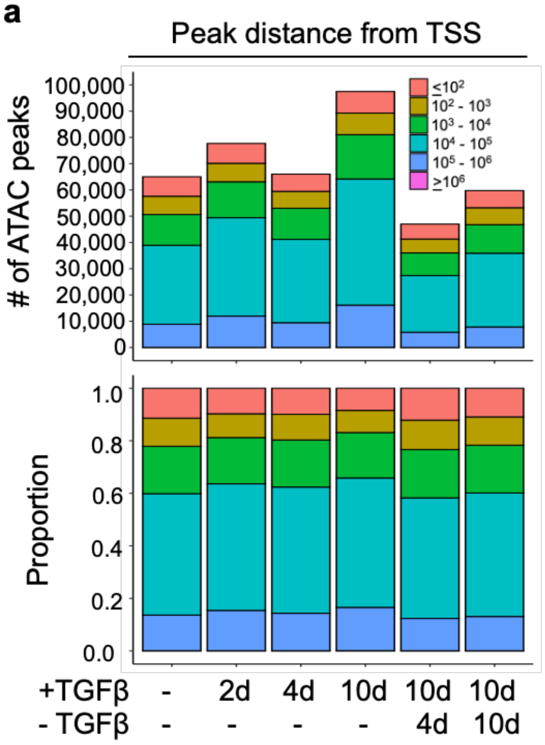


### Table 3

| Treatment | Max bp Distance from TSS | Min bp Distance from TSS | Mean bp distance from TSS | Median bp distance from TSS |
|---|---|---|---|---|
| Untreated | 1563344 | 0 | 49410 | 17558 |
| 2d +T | 1563342 | 0 | 54256 | 20912 |
| 4d +T | 1563344 | 0 | 51299 | 19533 |
| 10d +T | 1643527 | 0 | 58249 | 22834 |
| 10d +T, 4d -T | 1563348 | 0 | 45223 | 16080 |
| 10d +T, 10d -T | 1563331 | 0 | 47639 | 17521 |

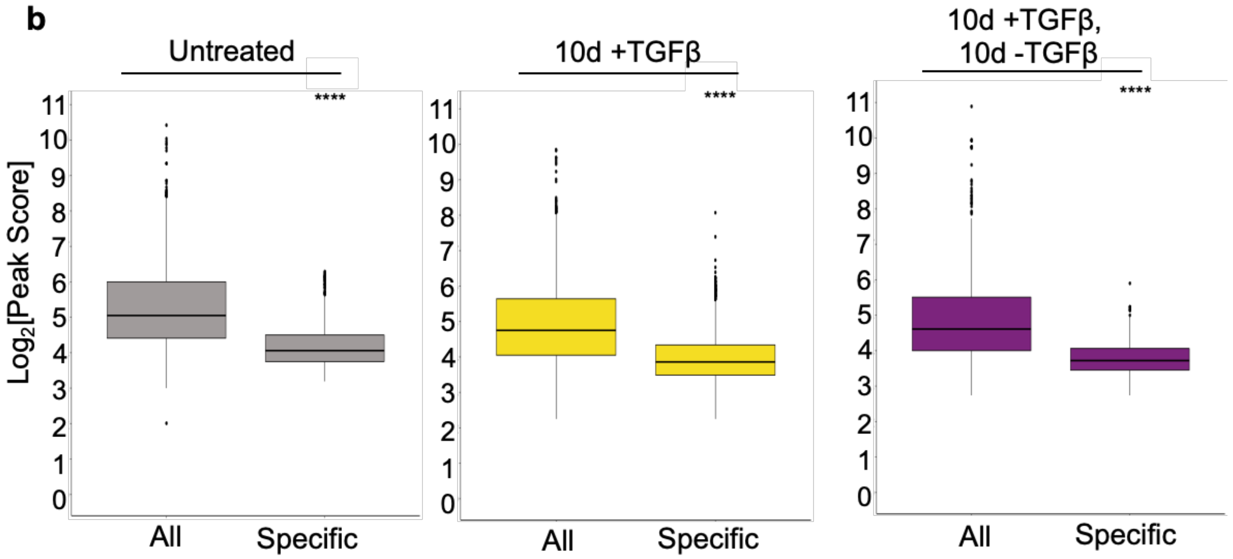


**Supplementary Figure 3.** Characteristics of ATAC peaks in long-term TGFβ treatment model. **a** Peak distance to closest transcription start site (TSS) of accessible regions of peaks common among replicates in long-term TGFβ-induced model. **b** $\log_2$[Peak Score] distribution for all and differential ($\log_2$[Differential Peak Score] $\geq$ 2) in untreated, 10d +TGFβ, and 10d +TGFβ, 10d -TGFβ conditions. Student's t-test used for statistical analysis. **** denotes $p \leq 0.0001$. **Table 3** Table representing maximum, minimum, mean, and median bp distance from TSS of accessible regions of peaks common among replicates in long-term model.

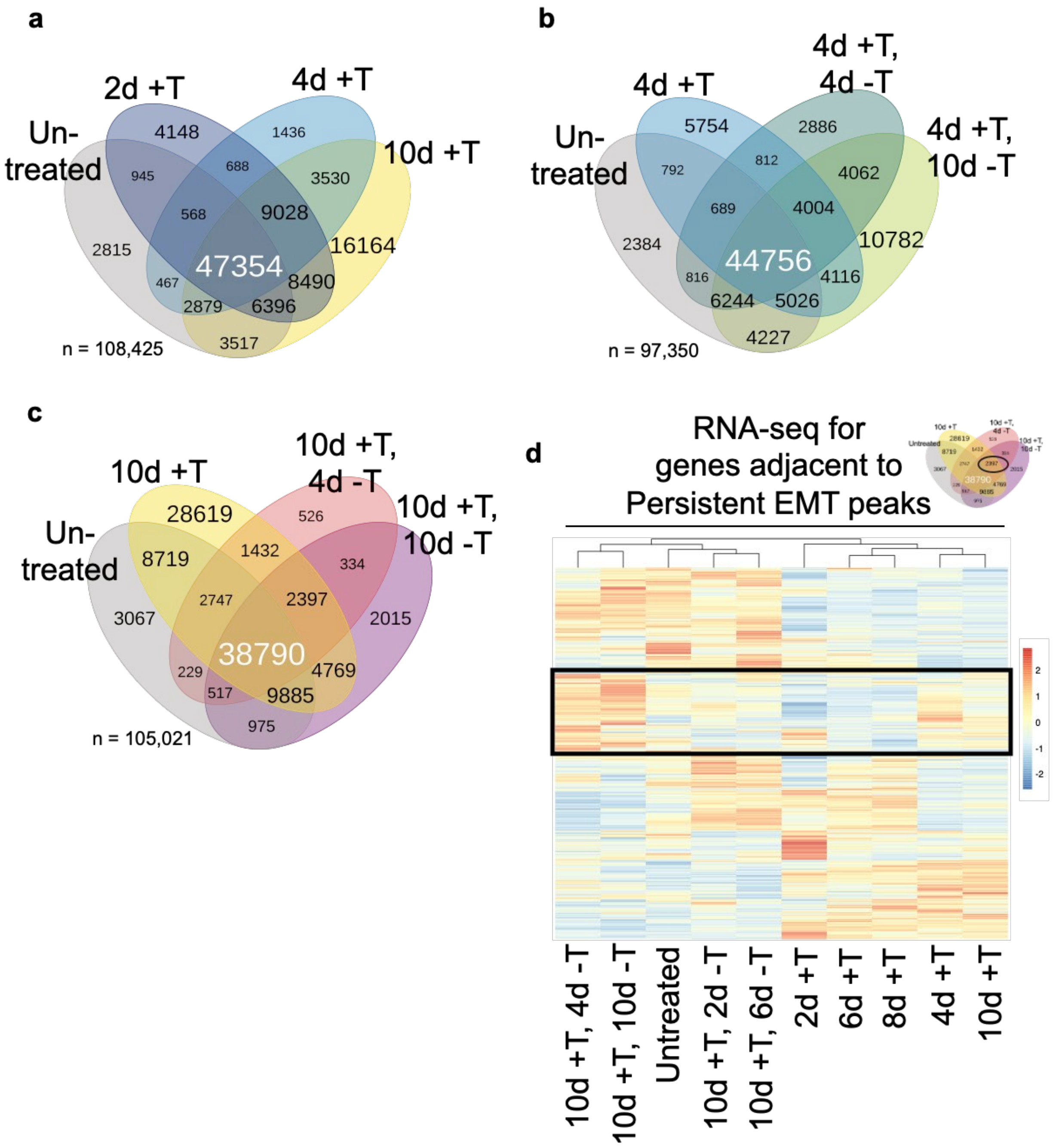


**Supplementary Figure 4.** Enumeration of peaks common to indicated TGFβ-treated and withdrawn treatments. **a-c** Venn diagram representing the overlap between ATAC peaks common among replicates in **a** untreated and TGFβ-induced conditions **b** short-term EMT model, and **c** long-term EMT model. **d** Unsupervised hierarchical clustering analysis of average mRNA expression by RNA-seq of genes annotated to ATAC peaks with persistent chromatin alteration following TGFβ treatment in long-term TGFβ-induced EMT model (n = 2,397).

**Table 4**

| | Hallmark Gene Sets |
|---|---|
| 1 | UV Response DN |
| 2 | Estrogen Response Early |
| 3 | Epithelial mesenchymal transition |
| 4 | TNFα Signaling via NFκB |
| 5 | Apoptosis |
| 6 | IL2 STAT5 Signaling |
| 7 | Hypoxia |
| 8 | Mitotic Spindle |
| 9 | TGFβ Signaling |
| 10 | Androgen Response |
| 11 | Inflammatory Response |
| 12 | Hedgehog Signaling |
| 13 | Apical Junction |
| 14 | p53 Pathway |
| 15 | Xenobiotic Metabolism |
| 16 | KRas Signaling Up |

**Table 5**

| | Breast and Mammary-Specific Gene Sets | Study |
|---|---|---|
| A | Mammary Stem Cell Up | Lim et al., 2010 |
| B | Luminal vs Mesenchymal Down | Charafe et al., 2006 |
| C | Luminal vs Basal Down | Charafe et al., 2006 |
| D | Pubertal Breast 4/5wk Up | McBryan et al., 2007 |
| E | Basal vs Luminal | Farmer et al., 2005 |
| F | Luminal B Down | Smid et al., 2008 |
| G | Pubertal Breast 3/4wk Up | McBryan et al., 2007 |
| H | Basal Up | Smid et al., 2008 |
| I | Invasive Breast Cancer Down | Poola et al., 2005 |
| J | Breast Cancer Relapse in Bone Down | Smid et al., 2008 |
| K | Ductal Invasive Up | Schuetz et al., 2006 |
| O | Basal Down | Smid et al., 2008 |
| L | Medullary vs Ductal Breast Cancer Down | Bertucci et al., 2006 |
| M | Luminal Mature Down | Lim et al., 2010 |
| N | Ductal Invasive Down | Schuetz et al., 2006 |
| P | ESR1 Laser Up | Yang et al., 2006 |
| Q | Copy Number Down | Climent et al., 2007 |
| R | Luminal vs Basal Up | Charafe et al., 2006 |

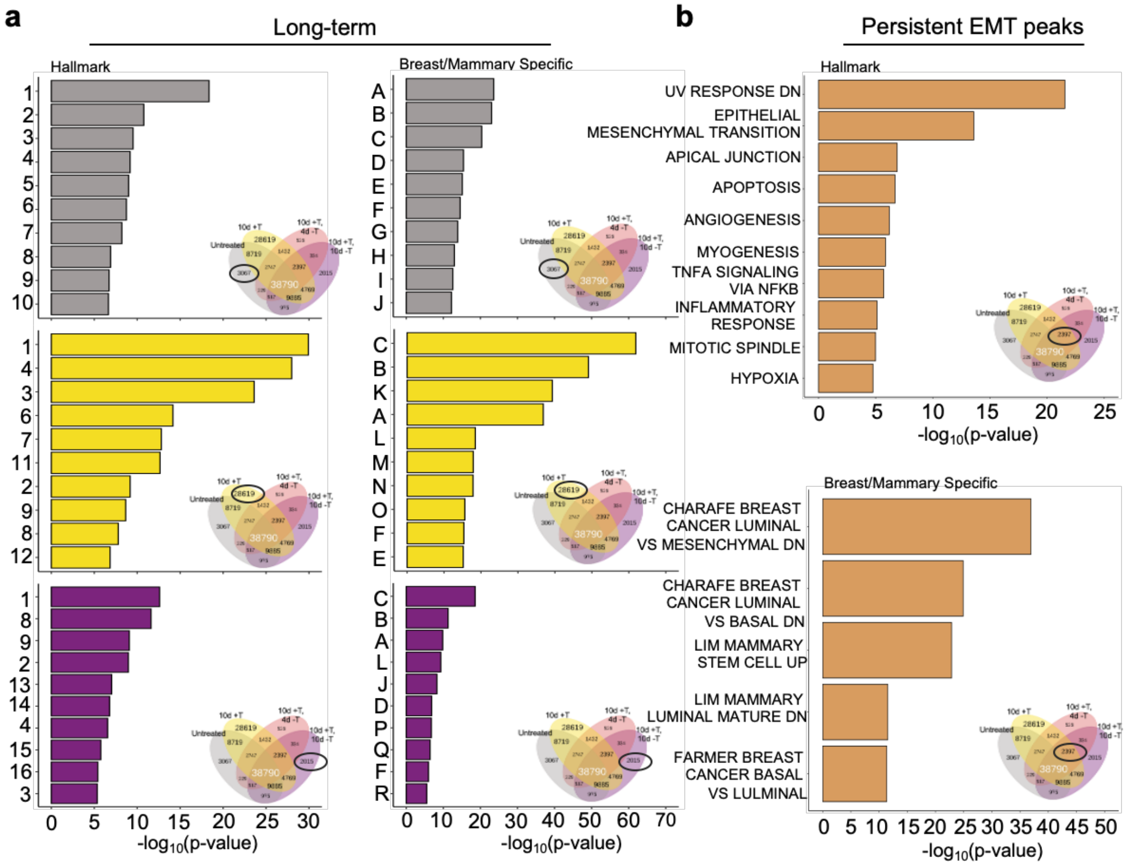


**Supplementary Figure 5.** EMT, mammary basal cell, and stemness gene sets are enriched in TGFβ induced and persistent peaks. **Table 4** Number key for GSEA msigdb hits for Hallmark gene sets. **Table 5** Letter key for GSEA msigdb hits for breast and mammary-specific gene sets. **a** GSEA msigdb hits (based on number key in **Table 4**) for ATAC peaks in Hallmarks gene sets (left) and hits (based on letter key in **Table 5**) for breast and mammary-specific gene sets (right) in long-term TGFβ-induced and –withdrawn conditions. Gray = untreated; yellow = 10d +TGFβ; purple = 10d +TGFβ, 10d -TGFβ. **b** GSEA msigdb hits for genes annotated to ATAC peaks with persistent chromatin alterations following TGFβ treatment and withdrawal in Hallmarks gene sets (top) and breast and mammary-specific gene sets (bottom).

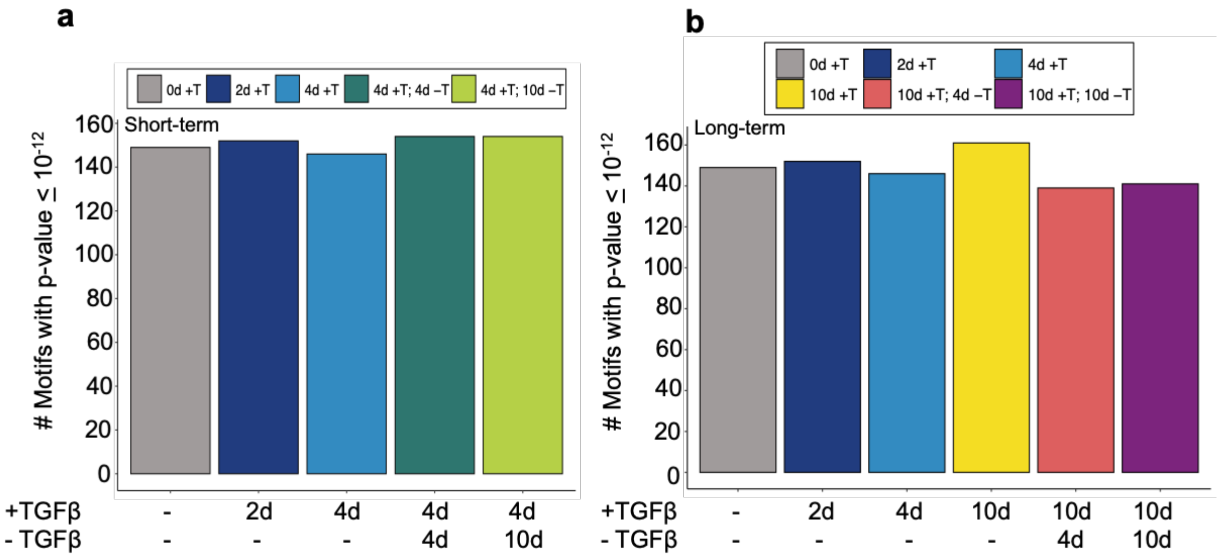


**Supplementary Figure 6.** Reversible EMT is driven by motifs utilized in baseline untreated conditions. Number of motifs considered highly-significant (p-value $\leq 10^{-12}$) in **a** short-term, and **b** long-term EMT models.

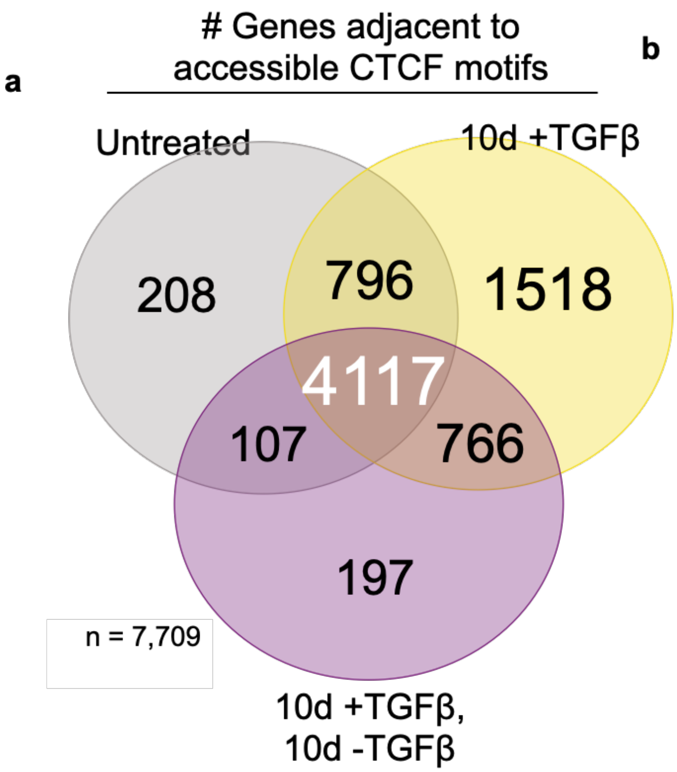


**Table 6**

| | Gene Set |
|---|---|
| 1 | CCANNAGRKGGC UNKNOWN |
| 2 | MEISSNER BRAIN HCP WITH H3K4ME3 AND H3K27ME3 |
| 3 | CAGGTG V$E12 Q6 |
| 4 | GO NEURON PART |
| 5 | GO REGULATION OF INTRACELLULAR SIGNAL TRANSDUCTION |
| 6 | GO NEUROGENESIS |
| 7 | ONDER CDH1 TARGETS 2 DN |
| 8 | MEISSNER NPC HCP WITH H3K4ME2 |
| 9 | GO POSITIVE REGULATION OF CELL COMMUNICATION |
| 10 | KOINUMA TARGETS OF SMAD2 OR SMAD3 |
| 11 | BENPORATH ES WITH H3K27ME3 |
| 12 | GO CELL DEVELOPMENT |
| 13 | GO CELL PROJECTION |
| 14 | GO REGULATION OF CELL PROJECTION ORGANIZATION |
| 15 | PEREZ TP53 TARGETS |
| 16 | GO RESPONSE TO OXYGEN CONTAINING COMPOUND |
| 17 | chr1q32 |

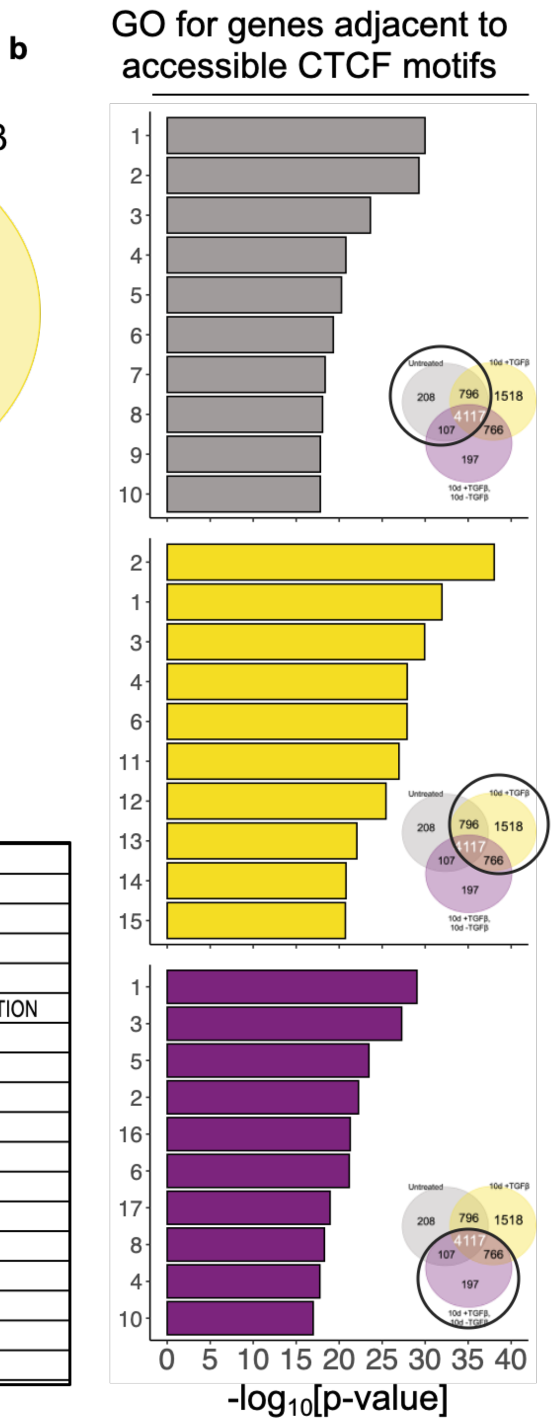


**Supplementary Figure 7.** Genes near engaged CTCF motifs are enriched for bivalent, neuronal, and cell projection-related gene sets. **a** Venn diagram representing the number of genes nearby accessible regions containing CTCF motifs for the indicated timepoints. **Table 6** GSEA msigDB gene sets determined to be highly-enriched among treatment conditions and their number key (first column). **b** GSEA msigDB enrichment (based on number key in **Table 6**) for top-10 enriched gene sets by condition. Gray = untreated; yellow = 10d +TGFβ; purple = 10d +TGFβ, 10d -TGFβ).
